# Parameter tuning differentiates granule cell subtypes enriching the repertoire of retransmission properties at the cerebellum input stage

**DOI:** 10.1101/638247

**Authors:** Stefano Masoli, Marialuisa Tognolina, Umberto Laforenza, Francesco Moccia, Egidio D’Angelo

## Abstract

The cerebellar granule cells (GrCs) form an anatomically homogeneous neuronal population which, in its canonical description, discharges regularly without adaptation. We show here that GrCs in fact generate diverse response patterns to current injection and synaptic activation, ranging from adaptation to acceleration of firing. Adaptation was predicted by parameter optimization in detailed GrC computational models based on the available knowledge on GrC ionic channels. The models also predicted that acceleration required the involvement of additional mechanisms. We found that yet unrecognized TRPM4 currents in accelerating GrCs could specifically account for firing acceleration. Moreover, adapting GrCs were better in transmitting high-frequency mossy fiber (MF) bursts over a background discharge than accelerating GrCs. This implied that different electroresponsive patterns corresponded to specific synaptic properties reflecting different neurotransmitter release probability. The correspondence of pre- and post-synaptic properties generated effective MF-GrC transmission channels, which could enrich the processing of input spike patterns and enhance spatio-temporal recoding at the cerebellar input stage.

## Introduction

The intrinsic variability in the ionic currents, the neuron’s morphology, and the neurotransmitter release dynamics are thought to be crucial for generating the richness of circuit properties and information-carrying capacity of brain microcircuits (Getting, 1989; Yarom and Hounsgaard, 2011; Gjorgjieva et al., 2016). A puzzling case is presented by cerebellar granule cells (GrC), the most numerous neurons of the brain (Herculano-Houzel, 2010). GrCs are located at the input stage of cerebellum, where they are thought to perform the fundamental operations of combinatorial expansion and spatio-temporal recoding predicted by the Motor Learning Theory (Marr, 1969; D’Angelo, 2016). While these operations would greatly benefit of a rich repertoire of signal transformation properties, GrCs appeared very homogeneous in size and shape since their original description (Golgi, 1906; Cajal, 1911). Later on, electrophysiological recordings from the cerebellar vermis of rodents reported a stereotyped firing pattern with spikes organized in regular discharges with little or no adaptation (D’Angelo et al., 1995; Brickley et al., 1996; Cathala et al., 2003). The experimental identification of a basic set of 8 ionic mechanisms (D’Angelo et al., 1998) allowed to model GrC firing in great detail (D’Angelo et al., 2001; Goldfarb et al., 2007; Diwakar et al., 2009; Dover et al., 2016) leading to a “canonical” description of GrCs but the question about the potential differentiation of firing properties remained open.

In a recent work (Masoli et al., 2017), automatic optimization procedures (Druckmann et al., 2007; Druckmann et al., 2011; Van Geit et al., 2016; Migliore et al., 2018) were applied to GrC models yielding a family of solutions with maximum ionic conductance values falling within the range of physiological variability. During short current injections (500-800 ms, as used in previous experiments), all GrC models conformed to the canonical firing pattern but, unexpectedly, prolonged current injections (2 sec) revealed a rich repertoire of adaptation properties. Here this prediction was tested experimentally by whole-cell recordings, which indeed revealed various degrees of firing adaptation and, in addition, also firing acceleration in some GrCs. This observation not just supported that conductance tuning would allow the emergence of a richness of electroresponsive properties but also implied the presence of yet unrecognized ionic mechanism causing firing acceleration.

Among the possible mechanisms causing firing acceleration there are transient receptor potential (TRP) channels (Subramaniyam et al., 2014; Stokum et al., 2018), which generate delayed depolarizing currents following intense firing and consequent activation of intracellular cascades (Nilius et al., 2005). Indeed, we have been able to measure TRP Melastatin 4 (TRPM4) currents in accelerating GrCs. These results suggest that GrC complexity in the cerebellar vermis is higher than previously thought, raising the number of ionic conductances required to determine the firing pattern. It was already known that GrC of vestibulo-cerebellum are specialized to slowdown firing modulation based on the expression of low-threshold calcium channels (Heath et al., 2014). Therefore, despite their morphological homogeneity, GrCs have differentiated conductance tuning and ionic channel expression, which could be further modified by fine variants in dendritic/axonal organization (Houston et al., 2017).

On a different scale, mossy fibers (MFs) convey to GrCs combinations of burst and protracted frequency-modulated discharges (Kase et al., 1980; van Kan et al., 1993; Chadderton et al., 2004; Jörntell and Ekerot, 2006; Rancz et al., 2007; Arenz et al., 2008) that are dynamically retransmitted at the MF-GrC synapses exploiting short-term plasticity mechanisms fine-tuned by vesicle release probability (*p*) (Mitchell and Silver, 2000b, a; Sola et al., 2004; Saviane and Silver, 2006; D’Errico et al., 2009). Here we observed that adapting GrCs exploit low-*p* synapses attaining a much higher signal to noise ratio (S/N) that accelerating GrCs.

These results show that, following Getting’s (1986) predictions, parameter variability at different scales supports the emergence of a richness of properties of potential physiological relevance in cerebellar GrCs. Tuning of adaption/acceleration and short-term plasticity generated a rich repertoire of filtering properties (Dean and Porrill, 2014; Rössert et al., 2015), which could substantially contribute to spatio-temporal recoding of synaptic input patterns at the cerebellum input stage (Marr, 1969).

## METHODS

### Experimental methods

All experimental protocols were conducted in accordance with international guidelines from the European Union Directive 2010/63/EU on the ethical use of animals and were approved by the ethical committee of Italian Ministry of Health (639.2017-PR; 7/2017-PR).

#### Slice preparation and solutions

Cerebellar GrCs were recorded from the vermis central lobe of acute parasagittal cerebellar slices (230 μm thick) obtained from 18- to 24-day-old Wistar rats of either sex. Slice preparation and patch-clamp recordings were performed as reported previously (Rossi et al., 1994; D’Angelo et al., 1995, 1997, 1998; Rossi et al., 1998; D’Angelo et al., 1999). Briefly, rats were decapitated after deep anesthesia with halothane (Sigma, St. Louis, MO), the cerebellum was gently removed and the vermis was isolated, fixed on a vibroslicer’s stage (Leica VT1200S) with cyano-acrylic glue and immersed in cold (2-3°C) oxygenated Kreb’s solution containing (mM): 120 NaCl, 2 KCl, 2 CaCl_2_, 1.2 MgSO_4_, 1.18 KH_2_PO_4_, 26 NaHCO_3_, and 11 glucose, equilibrated with 95% O_2_-5% CO_2_ (pH 7.4). Slices were allowed to recover at room temperature for at least 40 min before being transferred to a recording chamber mounted on the stage of an upright microscope (Zeiss, Germany). The slices were perfused with oxygenated Krebs solution (2 mL/min) and maintained at 32°C with a Peltier feedback device (TC-324B, Warner Instrument Corp., Hamden, CT, USA).

#### Patch-clamp recordings and data analysis

Whole-cell patch-clamp recordings from cerebellar GrCs (n=63) were performed with Multiclamp 700B [-3dB; cutoff frequency (fc), 10 kHz], sampled with Digidata 1440A/1550 interface, and analyzed off-line with pClamp10 software (Molecular Devices, CA, USA), MS Excel, Matlab (Mathworks, Natick, MA) and OriginPro software.

Patch-clamp pipettes were pulled from borosilicate glass capillaries (Hilgenberg, Malsfeld, Germany) and had a resistance of 7-9 MΩ before seal formation when filled with the intracellular solution containing (in mM): 126 potassium gluconate, 4 NaCl, 5 Hepes, 15 glucose, 1 MgSO_4_.7H_2_O, 0.1 BAPTA-free, 0.05 BAPTA-Ca^2+^, 3 Mg^2+^-ATP, 0.1 Na^+^-GTP, pH 7.2 adjusted with KOH. The calcium buffer is estimated to maintain free calcium concentration around 100 nM. Just after obtaining the cell-attached configuration, electrode capacitance was carefully cancelled to allow for electronic compensation of pipette charging during subsequent current-clamp recordings. The stability of whole-cell recordings can be influenced by modification of series resistance (R_s_). To ensure that R_s_ remained stable during recordings, passive cellular parameters were extracted in voltage-clamp mode by analyzing current relaxation induced by a 10 mV step from a holding potential of −70 mV. The transients were reliably fitted with a bi-exponential function yielding membrane capacitance (C_m_) of 3.1 ± 0.1 pF, membrane resistance (R_m_) of 1.1 ± 0.1 GΩ, and series resistance (R_s_) of 22.8 ± 1.1 MΩ. The −3 dB cell plus electrode cutoff frequency, f_VC_ = (2 R_s_C_m_)^−1^, was 2.6 ± 0.1 kHz (n = 63) (Silver et al., 1992; D’Angelo et al., 1993; D’Angelo et al., 1995).

GrCs intrinsic excitability was investigated in current-clamp mode by setting resting membrane potential at −65 mV and injecting 2 s current steps (from - 8 to 22 pA in 2 pA increment). The action potentials frequency was computed in two time windows of the duration of 500 ms, (0-500 ms and 1500-2000 ms respectively). In a subset of experiments (n=26) the MF bundle was electrically stimulated to investigate differences in synaptic transmission. The stimulation was performed with a large-tip (10-20 μm) patch-pipette filled with extracellular Kreb’s solution, via a stimulus isolation unit. The stimulation protocol comprised 1 sec of background stimulation (at either 5, 10, 20, 40, 60, 80 Hz) followed by 250 ms at 100 Hz burst stimulation, and was repeated twice. In some experiments slices were bath-perfused with Krebs added with 9-Phenanthrol (Sigma-Aldrich).

Data are reported as mean ± SEM, and, unless otherwise indicated, statistical comparisons are done using paired and unpaired Student’s t test.

#### Immunofluorescence

Immunofluorescence of cerebellar slices was performed as described previously (Laforenza et al., 2013). Rats, anesthetized as above described, were perfused transcardially and postfixed overnight in 4% paraformaldehyde in PBS. The fixed cerebella were cryo-protected with 30% sucrose solution in PBS, embedded in OCT (Cryostat embedding medium, Killik, Bio-Optica), and stored at −80°C. 20-μm-thick cryosections were treated for antigen unmasking by boiling in a microwave for 10 min in 0.01 M citrate buffer, pH 6. Slides were blocked with 3% BSA in PBS (blocking solution) for 30 min at room temperature. Successively, sections were incubated for 2 h at room temperature with the following affinity pure primary antibodies: rabbit polyclonal anti-TRPM4 (Cat.# ABN418; Merck Millipore, Darmstadt, Germany) and mouse monoclonal anti-Aquaporin 4 antibody [4/18] (ab9512; Abcam, Cambridge, UK), diluted 1:60 and 1:100 in blocking solution, respectively. Slices were washed 3 times and then stained simultaneously with two secondary antibodies: goat anti-mouse IgG H&L (Alexa Fluor® 488) preadsorbed (1:400 diluition; ab150117; Abcam, Cambridge, UK) and rhodamine red -X-coniugated affinity pure goat anti-rabbit IgG (H+L) (1: 50 dilution; 111-295-003; Jackson ImmunoResearch Europe Ltd, Cambridge, UK) for 1 h at room temperature. Slides were then washed 3 x 5 min with PBS, mounted in BrightMount/Plus aqueous mounting medium (ab103748; Abcam, Cambridge, UK). Slides were examined with a TCS SP5 II confocal microscopy system (LeicaMicrosystems) equipped with a DM IRBE inverted microscope (LeicaMicrosystems). Images were acquired with 20, 40, or 63 objectives and visualized by LAS AF Lite software (Leica Microsystems Application Suite Advanced Fluorescence Lite version 2.6.0). Negative controls were performed by incubating slices with non immune serum.

### Computational modeling

The GrC model used in this study was written in Python 2.7/NEURON 7.6 (Hines and Carnevale, 2001; Hines et al., 2009). The model derived from previous ones (D’Angelo et al., 2001; Nieus et al., 2006; Diwakar et al., 2009) and was upgraded to account for advanced mechanisms of spike generation and conduction in the axon (Dover et al., 2016). The ionic channels were distributed among dendrites, soma, hillock, axonal initial segment (AIS), ascending axon (AA) and parallel fibers (PF) (Masoli et al., 2017). The maximum ionic conductances (G_max_) were optimized using routines based on genetic algorithms (BluePyOpt) (Deb et al., 2002; Zitzler and Künzli, 2004; Van Geit et al., 2016; Masoli et al., 2017). The optimization was run iteratively to improve models fitness to an experimental “template” through the automatic evaluation of “feature” values parameterizing the spike properties of the template. Out of 3 rounds of optimization we obtained more than 600 GrC models, 150 of which were chosen randomly for further analysis. The simulation workflow was fully automated and parallelized and allowed to perform a series of protocols for: 1) parameter optimization, 2) simulation of the last generation, 3) filtering of the population based on the experimental properties, 4) simulation of each individual with protocols identical to those used experimentally.

The electrotonic structure of the granule cell models was derived from (Dover et al., 2016) along with passive parameters (Table 1 in Supplemental Material). The GrC ionic channel models and distributions were taken from previous papers and updated according to the latest literature when needed (Table 2 in Supplemental Material). The model included Na^2+^ channels (Nav1.6 with and without FHF), K^+^ channels (Kv1.1, Kv1.5, Kv2, Kv4.3, Kv3.4, Kv7), Ca^2+^ channels (Cav2.2) and the newly reported TRPM4 channels, along with a Calretinin based calcium buffer (Gall et al., 2003). Synaptic transmission was modelled using the Tsodyks and Markram scheme (Tsodyks et al., 1998) adapted as in (Nieus et al., 2006).

#### Feature extraction

GrC discharge features were extracted from experimental traces of GrCs showing regular firing. The data were taken at three current injection steps (10, 16 and 22 pA), using the “Electrophys Feature Extraction Library” (EFel) (https://github.com/BlueBrain/eFEL) (Van Geit, 2015). Accordingly to (Masoli et al., 2017), the features comprised resting membrane potential, spike width and height, fast and slow AHP depth, mean spike frequency, time-to-first spike, adaptation and coefficient of variation of the interspike interval (ISI-CV) (Table 3 in Supplemental Material).

#### Model optimization and validation

Automatic optimization of maximum ionic conductances was performed using the “Blue Brain Python Optimization Library” (BluePyOpt) (Van Geit et al., 2016), which is written in Python and C and uses the IBEA algorithm. Each optimization had an initial population of 288 individual, yielding a 576 final population and was performed for 12 generation, with fixed time step, at a temperature of 32°. A single optimization took about 5 hours to be completed. After each optimization, the best individuals of the last generation were used as a guidance to reshape the ionic conductance ranges and to improve the fitness values. The parameter range for maximum ionic conductances was limited by physiological measurements as in (Masoli et al., 2017). The results of optimization were validated by evaluating spike generation and conduction. A model was discarded when (1) spike generation in the AIS failed, (2) spike conduction speed or spike amplitude in the AA and PFs was decremental, (3) spike frequency was different in soma and axon (>± 1 spike/sec) (e.g. see (Dover et al., 2016)).

#### Model simulations and analysis

The BluePyOpt template was customized to allow the simulation and validation of each optimized model and to improve the simulation speed by using a Python module providing access to the Message Passing Interface (MPI4py). The ionic channel conductances were uploaded from the final GrC population file. The simulations were run for 2.2 s at 32° with fixed time step (Hines and Carnevale, 2008) and with the same experimental current injections (10, 16 and 22 pA). The voltage traces were recorded and saved, for each model, from four locations: soma, AIS, final section of the AA and final section of PFs. Optimizations and simulations were carried out using 8 nodes (36 cores each) of the “HBP Blue Brain 5” cluster (BB5), located at the CSCS facility in Lugano, while coding, testing, debugging and additional simulations were carried out on an 8core/16 threads CPU (AMD Ryzen 1800x with 32GB of ram). The simulations were analyzed by the same routines and statistical tests used for the experimental data.

#### Public distribution of models

The optimization Python notebook will be made available as a use case on the Brain Simulation Platform (BSP) along with the BluePyOpt version of GrC models (Masoli et al., 2017). Validated models will be made available to be directly simulated, on the same platform, using BlueNaaS (BBP - Neuron as a service). The experimental data, simulations and optimization results will be made available in the form of a “live paper” on the BSP.

## RESULTS

The cerebellar GrCs are thought to form a highly homogeneous neuronal population, which is silent at rest and then responds to current injection with regular repetitive firing showing little or no adaptation. However, the generation of GrC model families using automatic parameter optimization (Masoli et al., 2017) predicted that, following the initial 500-800 ms discharge usually considered for the assessment of neuronal firing, the GrCs may show adaptation to various degrees.

### Different intrinsic electroresponsive properties in cerebellar GrCs

In order to accurately assess the GrC firing properties, whole-cell recordings were carried out in current-clamp configuration while delivering 2-sec current steps at different intensities from the holding potential of −65 mV. This protocol differs from those used previously (D’Angelo et al., 1995; Brickley et al., 1996; D’Angelo et al., 1998; Cathala et al., 2003) simply because it is longer than usual. All recorded GrCs were silent at rest and their response frequency at 500 ms showed little or no adaptation (Fig. 1A). However, surprisingly enough, while initial GrC responses corresponded to the classical description, at longer times they showed a richness of different properties (Fig. 1A). In some cells firing remained stable (*non-adapting*), in others it slowed-down or even stopped (*adapting*), while yet in others it increased (*accelerating*). While changes in the initial 500 ms were less than about ±20%, changes over the whole 2000 ms time-window could exceed ±80% (Fig. 1B).

The cerebellar GrCs normally show a linear firing frequency increase with current injection (D’Angelo et al., 1995; Brickley et al., 1996; D’Angelo et al., 1998; Rossi et al., 1998; Cathala et al., 2003). During the first 500 ms of the response, both in non-adapting, adapting and accelerating GrCs, the firing frequency increased monotonically and almost linearly with current injection (*f*_*initial*_ /*I* plots) (Fig. 1C).

In summary, when observed during the first 500 ms of the response to current injection, GrC firing frequency was almost stable and the input-output relationships linear, conforming to general knowledge. However, a richness of electroresponsive properties emerged at longer response times (here up to 2000 ms).

The firing frequency changes occurring over 2000 ms current steps were assessed by calculating the *intrinsic frequency change IFC* = [(*f*_*final*_−*f*_*initial*_)/*f*_*initial*_]%, in which *f*_*initial*_ and *f*_*final*_ are the spike frequencies at the beginning and end of current injection. In this equation, adaptation and acceleration are characterized by IFC<0 and IFC>0, respectively, while *IFC*=0 occurs in the absence of changes. Characteristically, the accelerating GrC showed a IFC positive peak around 10 pA current injection, while the other GrCs showed negative IFC values (see Fig. 1D).

**Figure 1.**
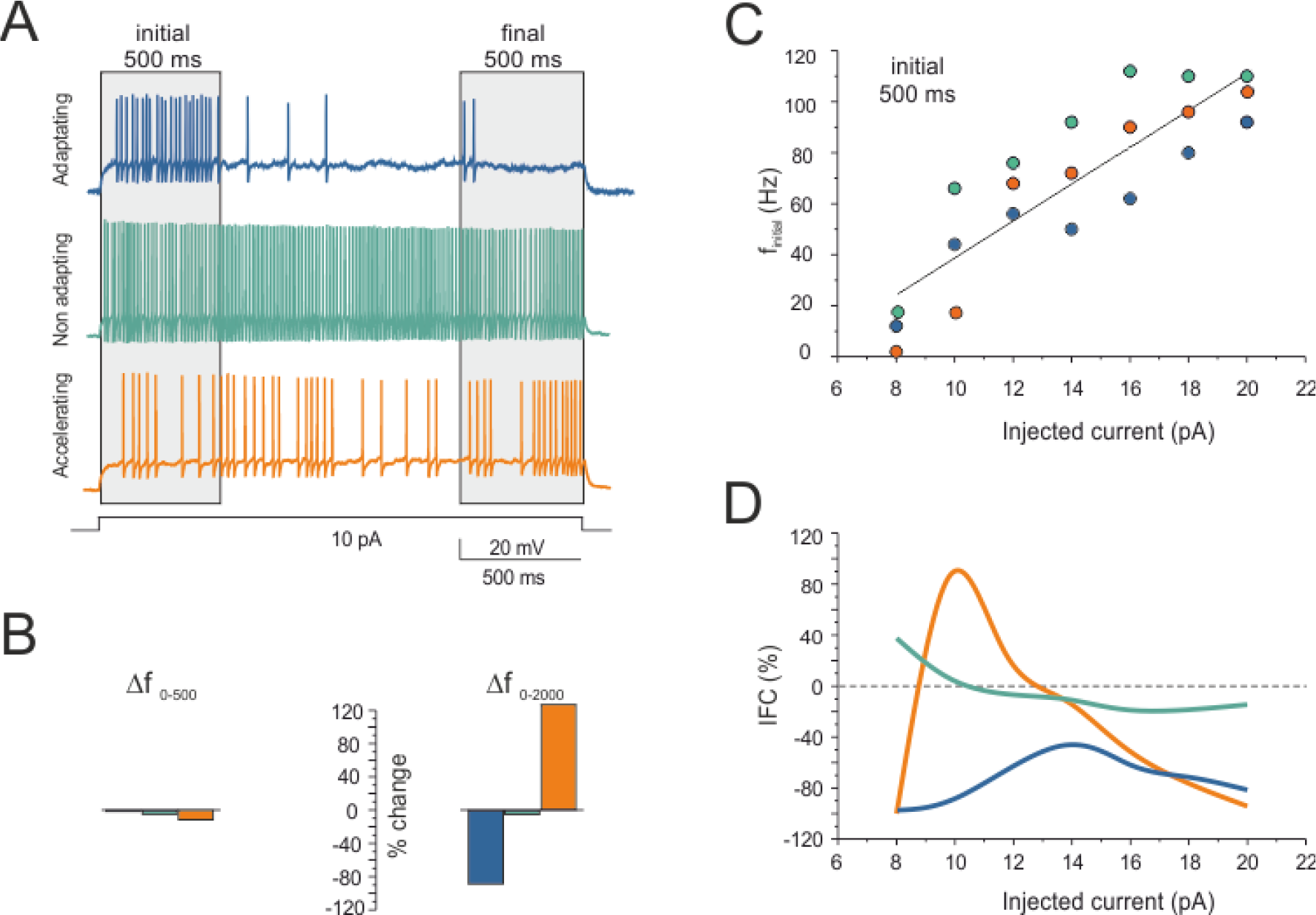
Different GrC properties: adaptation and acceleration. Three exemplar GrC recordings are shown, one *adapting*, one *non-adapting*, one *accelerating*. The same color codes are used consistently in the figure. **(A)** Voltage responses to 2000 ms - 10 pA current injection from the holding potential of −65 mV. Spike frequency initially remains stable in all the three cells but it shows different trends thereafter. **(B)** Δf _0-500_ and Δf _0-2000_ are the spike frequency % changes after 500 ms and 2000 ms, respectively. **(C)** In *f*_*initial*_ /*I* plots, spike frequency increases almost linearly with the injected current intensity in both the accelerating, adapting and non-adapting granule cells (compound R^2^=0.90, p<0.001). **(D)** Plot of the intrinsic frequency change IFC *vs.* injected current for the three GrCs. A positive peak is apparent in the accelerating GrC at 10 pA current injection, while negative IFC values prevail in the other GrCs.

### Different synaptic excitation properties at the MF-GrC relay

The different firing properties of GrCs could have an impact on retransmission of MF discharges. This issue was addressed by stimulating the MF bundle at frequencies *f*_*stim*_ = 5-100 Hz and measuring the GrC response frequency, *f*_*resp*_. GrCs showed different transmission properties, from one-to-one responses over the whole input frequency range to responses faster or slower than the input (Fig. 2A). These properties were analyzed using *SFC*/*f*_*stim*_ plots, where *SFC* = [*(f*_*resp*_ − *f*_*stim*_)/*f*_*stim*_]% is the *synaptic frequency change* (Fig. 2B). The *SFC*/*f*_*stim*_ plots showed distinctive trajectories (Fig. 2B). The accelerating GrCs showed *SFC* > 0 (enhanced output) around 20 Hz, while the other GrCs showed *SFC* < 0 all over the frequency range.

Since MF-GrC transmission involves high-frequency bursts (Chadderton et al., 2004), we evaluated the MF-GrC signal/noise ratio, S/N (Fig. 2C). This was measured when a *signal* (high-frequency burst at 100 Hz) was delivered over *noise* (background activity at 20 Hz) (see Fig. 2C). The adapting GrCs suppressed the background efficiently but allowed high frequency burst transmission resulting in high S/N. The non-adapting and accelerating GrCs showed less efficient background suppression resulting in lower S/N. Thus, GrCs with different adaptation / acceleration properties also showed differential filtering of MF activity.

In adapting and accelerating GrCs, a voltage-clamp protocol was run to monitor the effectiveness of synaptic stimulation. The average EPSC amplitude was −39.5±4.8 pA (n=24), corresponding to activation of ~2 synapses on average (e.g. cf. (Silver et al., 1996; Sola et al., 2004). The paired-pulse ratio (PPR) in 5-100 Hz trains was 0.88±0.05 in strong-adapting GrCs (n=4 cells, 47 measures) and 0.53±0.1 in accelerating GrCs (n=5 cells, 46 measures). By comparison with the precise determinations carried out on this same synapses (Sola et al., 2004), these estimates suggested that adapting GrCs were activated by synapses with lower release probability than accelerating GrCs (n=93, p<0.01, unpaired *t*-test).

**Figure 2.**
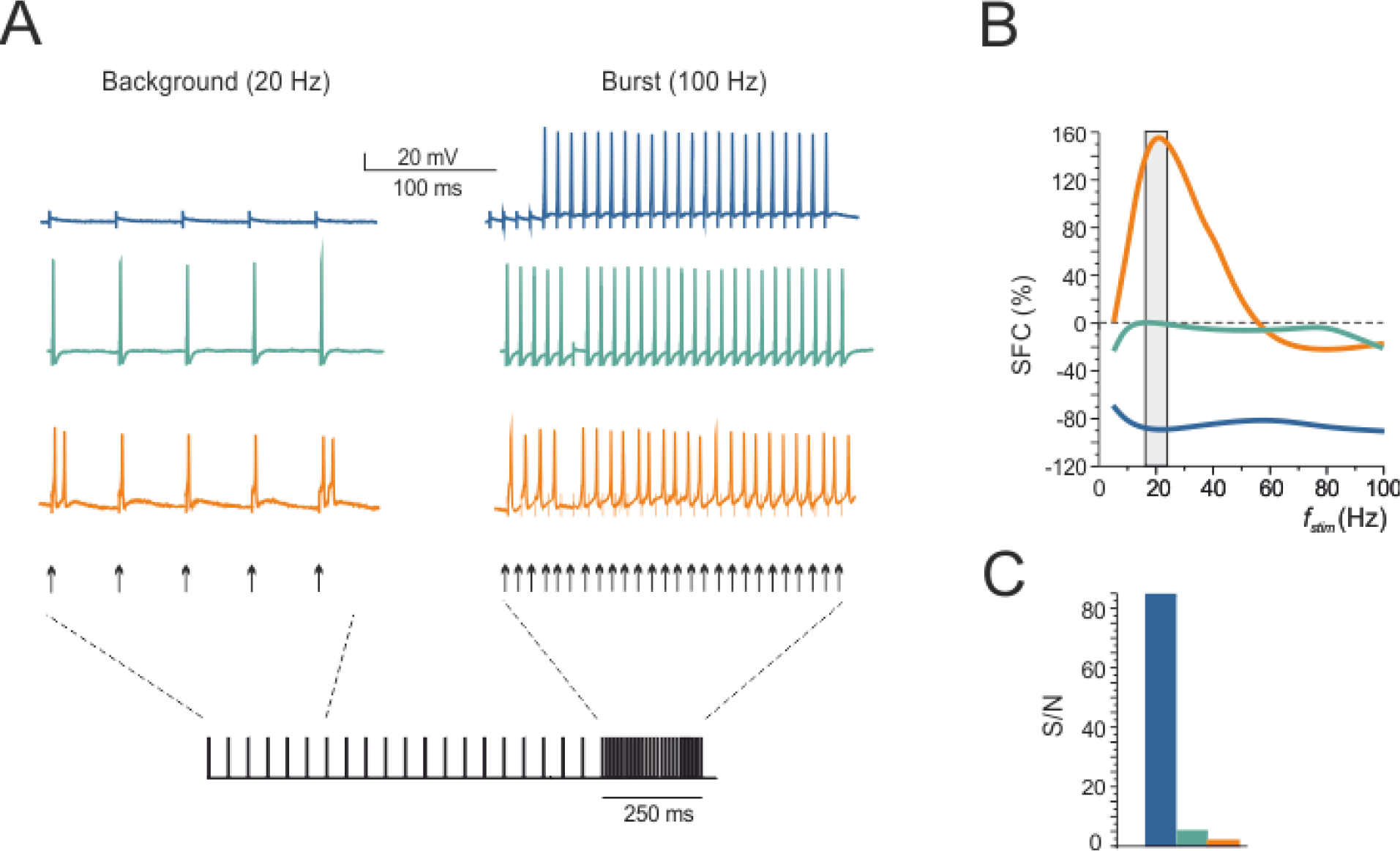
Different MF-GrC properties: synaptic responsiveness and S/N. Three exemplar granule cell recordings are shown, one *adapting*, one *non-adapting*, one *accelerating*, as defined by their intrinsic electroresponsiveness (same cells and color codes as in Fig. 1). **(A)** Voltage responses of exemplar GrCs to electrical stimulation of the MF bundle (1 sec continuous stimulation at 20 Hz followed by 250 ms at 100 Hz) from the holding potential of −65 mV. **(B)** Plot of the synaptic frequency change SFC *vs.* stimulus frequency for the three GrCs shown in A. A positive peak is apparent in the accelerating GrC at 20 Hz background stimulation, while negative SFC values prevail in the other GrCs. **(C)**. Signal-to-noise ratio, (S/N = f_resp_@100 Hz/f_resp_@20 Hz) for the three GrCs in A-B. Note that S/N is much higher in adapting that in the other two GrCs.

### Average properties of GrC subtypes

In order to determine whether the different GrC responses represented a continuum or rather could be grouped into subpopulations, an unbiased *k*-means cluster-analysis was applied to the whole population of recordings (n=63 at 10 pA) using *IFC* and *f*_*initial*_ as features (Fig. 3A). The *k*-means analysis was performed at low current intensity were differences among GrCs were more evident. The *k*-means analysis identified four statistically different data clusters corresponding to *strong-adapting* GrCs (n=23), *mild-adapting* GrCs (n=19), *non-adapting* GrCs (n=13) and *accelerating* GrCs (n=8) (Fig. 3A) (the adapting granule cells were actually subdivided into two groups). The cells identified in the four clusters were then used to analyze their average properties.

In the average *IFC*/*I* plot (Fig. 3B), non-adapting, mild-adapting and strong-adapting GrCs showed differential adaptation at low current injection (<14 pA) but converged toward a similar adaptation level at high current injection (20 pA). The accelerating GrCs showed increased intrinsic electroresponsiveness at low current injection (around 10 pA) but decreased it to the level of the other GrCs at higher current injections (20 pA).

In the average *SFC*/*I* plot (Fig. 3C), non-adapting, mild-adapting and strong-adapting GrCs showed differential response regimens at low synaptic stimulation frequencies (<50 Hz) but converged toward a similar response level at higher synaptic stimulation frequencies (100 Hz). The accelerating GrCs showed increased synaptic responsiveness at low input frequencies (around 20 Hz) but decreased it to the level of the other GrCs at higher input frequencies (100 Hz).

The average *IFC* and *SFC* plots showed similar trends for the four GrC categories. The correlation between *IFC* and *SFC* (Fig. 3D) was evaluated in a characteristic point corresponding to the peak of accelerating GrCs, i.e. using IFC at 10 pA and SFC at 20 Hz. The *IFC@10 pA/SFC@20 Hz* plot actually revealed a linear correlation (R^2^=0.70).

S/N was also evaluated at the cell population level in the four GrC clusters identified by k-means analysis (Fig. 3E). This analysis revealed that S/N was indeed progressively lower when passing from strong-adapting to mild-adapting, non-adapting and accelerating GrCs.

**Figure 3.**
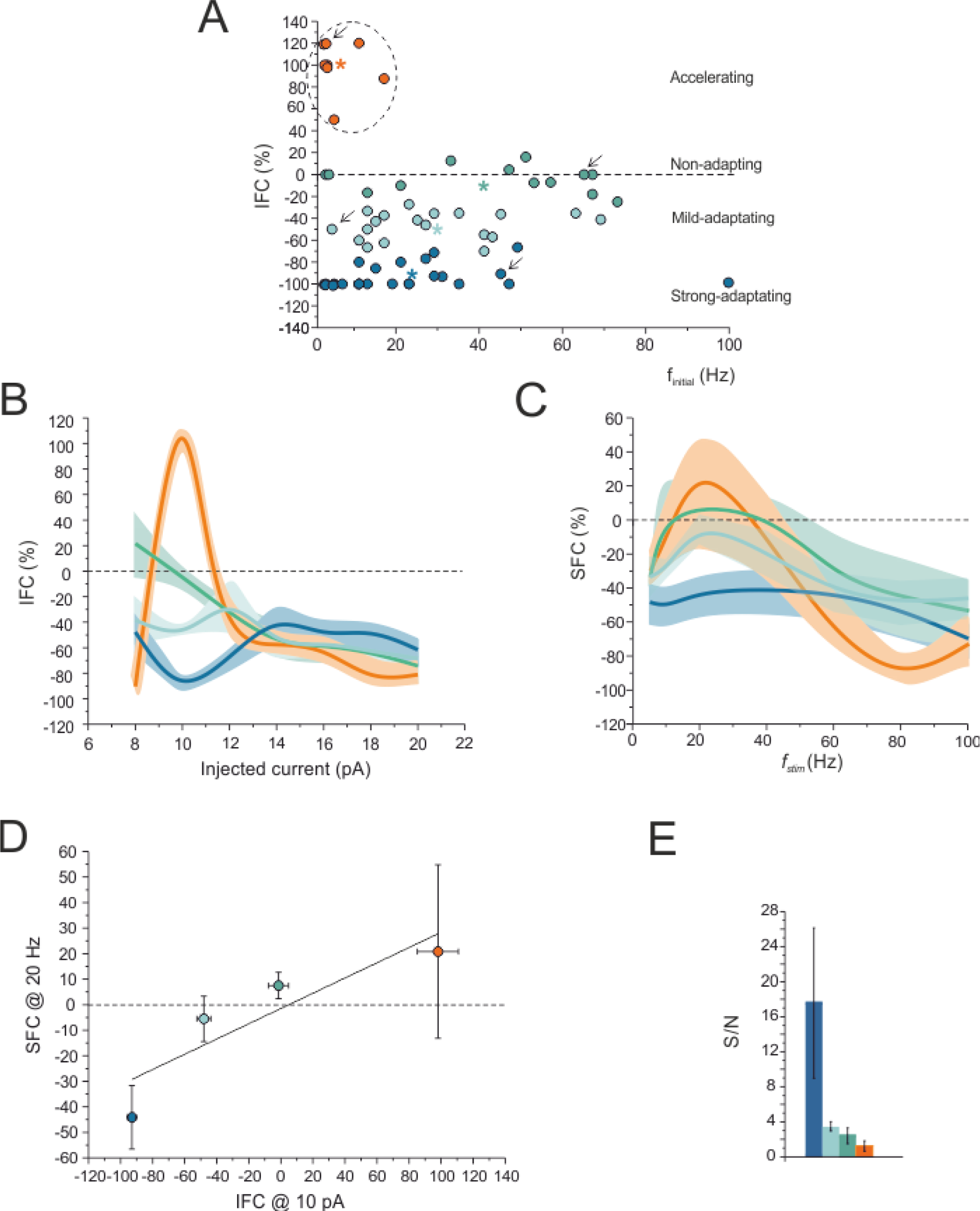
Average properties: granule cell subtypes. **(A)** *K*-means cluster-analysis applied to the whole population of recordings (n=63 at 10 pA) using *IFC* and *f*_*initial*_ as features. *K*-means identifies four statistically different data clusters (p =3.7e-12; Kruskal-Wallis test). The centroids of the 4 subpopulations are indicated by stars. **(B, C)** The cells identified in the four clusters were used to construct average *IFC*/*I* and *SFC*/*I* plots. Note, in both graphs, the similar and progressive change of properties from strong-adapting to mild-adapting, non-adapting and accelerating GrCs. **(D)** *IFC@10pA* and *SFC@20Hz* reveal a linear correlation (R^2^=0.70). **(E)** The cells identified in the four cluster are used to construct an average *S*/*N* histogram (S/N = f_resp_@100 Hz/f_resp_@20 Hz). Note the progressive decrease of S/N from adapting to accelerating GrCs. Color codes as in Fig. 1.

### TRP current expression in accelerating GrCs

The depolarization-induced slow current (DISC) is a depolarizing current gated by the raise of intracellular Ca^2+^ that follows action potential bursts and NMDA channel activation, and can typically generate a secondary burst after 1.5-2 secs like that observed in accelerating GrCs (Shin et al., 2008; Shin et al., 2009; Menigoz et al., 2016). DISC is typically generated by TRPM4 (Kim et al., 2013), a Ca^2+^-activated non-selective cation channel which provides a strong depolarizing drive upon Ca^2+^ entry (Petersen, 2002). TRPM4 currents were elicited in GrCs using the same voltage-clamp protocol adopted in PCs (Kim et al., 2013) (Fig. 4A). TRPM4-like currents could indeed been elicited in accelerating GrCs (n=4 out of 4). These currents occurred after 1780±210 ms and were typically burst-like with an average charge transfer of −26.00±4.21 pA*ms (n=4) (Fig. 4B). In order to confirm TRPM4 specificity, the preparations were perfused with 100 μM 9-Phenanthrol, a specific TRPM4 channel blocker (Mrejeru et al., 2011; Kim et al., 2013), which completely blocked the currents (n=4 out of 4). The spike discharge recorded after switching to current-clamp in these same accelerating GrCs showed increased frequency in correspondence to the TRP current, with *IFC@10 pA* = 86.1±27.7% (Fig. 4A). Conversely, non accelerating GrCs never showed the TRPM4-like current (n=12 out of 12).

The expression of TRPM4 channels in granule cells was verified by immunohistochemistry. Staining with anti-TRPM4 antibodies revealed immunofluorescence in the membrane of most GrCs (Fig. 4C). A recent investigation demonstrated that TRPM4 protein is abundantly expressed in cerebellar PCs, while its distribution in the granular layer was not evaluated (Kim et al., 2013). However, according to the *in situ* hybridization data from the Allen Institute of Brain Science (http://mouse.brain-map.org/), TRPM4 channels should also be expressed in GrCs. To validate this preliminary information at protein level, we carried out immunohistochemistry on the rat cerebellar granular layer by using a rabbit polyclonal antibody raised against the N-terminal domain of human TRPM4 (Kim et al., 2013). GrCs were identified by co-staining the slices with DAPI, as recently described in (Mapelli et al., 2017). Confocal microscopy revealed that TRPM4 protein was expressed in GrCs both on the plasma membrane and within the cytosol (Fig. 4C). This subcellular pattern of expression is similar to that reported in other brain areas (Teruyama et al., 2011; Lei et al., 2014). As expected, the antibodies against TRPM4 strongly stained also Purkinje cells (Kim et al., 2013). When the slices were labelled with DAPI, but not with the primary antibody, only background levels of immunoreactivity were detected, thereby confirming the antibody specificity (not shown).

**Figure 4.**
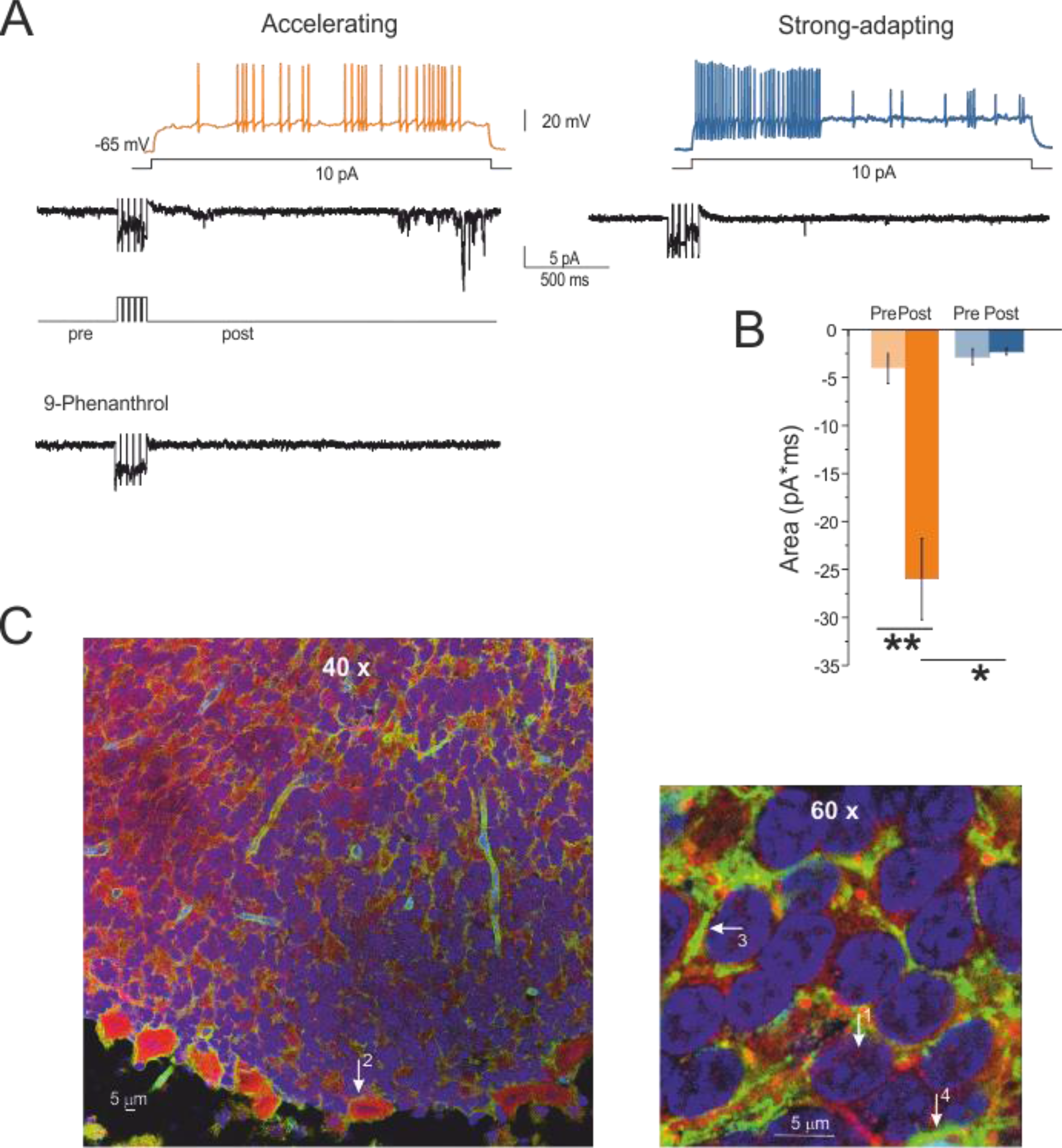
TRP channels in accelerating GrCs. **(A)** The voltage traces show the GrC response to a 10 pA step current in an accelerating GrC and in a strong-adapting GrC. A 5 steps depolarizing protocol is applied to the same two GrCs to uncover the depolarization-induced slow current, DISC. In the accelerating GrC, the current traces reveal a DISC, which typically appears as a delayed burst of rapid events. DISC disappears after perfusing a specific TRPM4 channel blocker, 9-Phenantrhol (100 μM). No DISC is visible in the strong-adapting GrC (all currents low pass filtered at 500 Hz). **(B)** The average area underlying the baseline shows a significant current increase after the application of the depolarizing protocol in accelerating GrCs (p<0.01, n=4, paired *t*-test) but not in strong-adapting granule cells. Stars indicate statistical significance of differences (* p<0.05, ** p<0.01). **(C)** Fluorescent images obtain by thin sections of the cerebellar vermis stained with antibodies against TRPM4 (red), aquaporin-4 (green), DAPI (blue). TRPM4 is expressed in the cell membrane surrounding the GrCs soma as well as in small grains in the cytoplasm (arrow 1). Aquaporin-4 stains glial sheets delimiting the GrC soma (arrow 3) and contours the blood vessels (arrow 4), supporting the specificity of TRPM4 staining for GrCs. DAPI stains the cell nuclei. TRPM4, aquaporin-4 and DAPI staining are also present in the molecular layer (TRPM4 are strongly expressed in PCs; arrow 2) (Kim et al., 2013). (Left 40x, right 63x objective).

### Computational modelling predicts parameter tuning in GrC subtypes

Since it is experimentally unpractical to determine the balance of multiple ionic conductances in single GrCs, we inferred membrane mechanisms from simulations using biophysically detailed data-driven models (Fig. 5A) (D’Angelo et al., 2001; Nieus et al., 2006; Diwakar et al., 2009; Dover et al., 2016; Masoli et al., 2017) incorporating multiple types of ionic channels on the dendrites, soma, hillock, axonal initial segment, ascending axon and parallel fibers (Fig. 5B; see also Supplemental Fig. 1). The hypothesis that the known set of ionic channels was indeed sufficient to explain the GrC firing subtypes was explored using automatic optimization of maximum ionic conductances (G_max_) (Deb et al., 2002; Zitzler and Künzli, 2004; Van Geit et al., 2016; Masoli et al., 2017) yielding a family of solutions that fit the experimental “template” (Fig. 5C).

The models were first optimized without TRPM4 channel coupling toward a template taken from the first 500 ms discharge in a non-adapting GrC. All the models could faithfully reproduce the stable regular firing behaviour typical of the first 500 ms of discharge (Fig. 5C) while, interestingly, adapting properties emerged at later times. Moreover, these GrC models clustered into three groups identifying the three subtypes observed experimentally (k-means analysis, Fig. 5D). Therefore, the ability to generate adaptation to various extents was intrinsic to the ionic channel complement through fine-tuning of G_max_ values (see Supplemental Fig. 2). Among the numerous ionic channels, the most systematic G_max_ change was that of Cav2.2 channels, that regularly degraded from strong-adapting to mild-adapting to non-adapting GrCs (Fig. 5E), suggesting that Cav2.2 were indeed the key regulators of adaptation (see below). The coupling of TRPM4 channels to Ca^2+^ through Calmodulin allowed to obtain accelerating GrCs (Nilius et al., 2005) (Fig. 5C). An exemplar case is reported in Fig. 6.

In the models, the MF - GrC synapse was implemented using a dynamic representation of the vesicle cycle, that could faithfully reproduce MF-GrC short-term plasticity including EPSC depression and facilitation (Fuhrmann et al., 2002; Nieus et al., 2006; Arleo et al., 2010; Solinas et al., 2010; Nieus et al., 2016). The different GrC models were stimulated synaptically with a protocol identical to that used for experimental recordings. An exemplar case is reported in Fig. 7. According to experimental estimates (see above and (Sola et al., 2004; Sgritta et al., 2017)), the GrC model was activated by 2 synapses with a higher release probability (p=0.5) for accelerating and non-adapting GrCs than for strong-adapting GrCs (p=0.1). The GrC model responses at different frequencies (Fig.7A), as well as the SFC/f_stim_ plots (Fig. 7B) and the S/N values (Fig. 7C), were remarkably similar to those obtained experimentally (cf. Fig. 2).

As a whole, GrC and MF models parameter tuning yielded electroresponsive and synaptic transmission properties that closely matched those observed experimentally (cf. Figs 6-7 to Figs 1-2). And since the models effectively captured the phenomenological properties of GrC subtypes, they were further used to infer the underlying ionic mechanisms.

**Figure 5.**
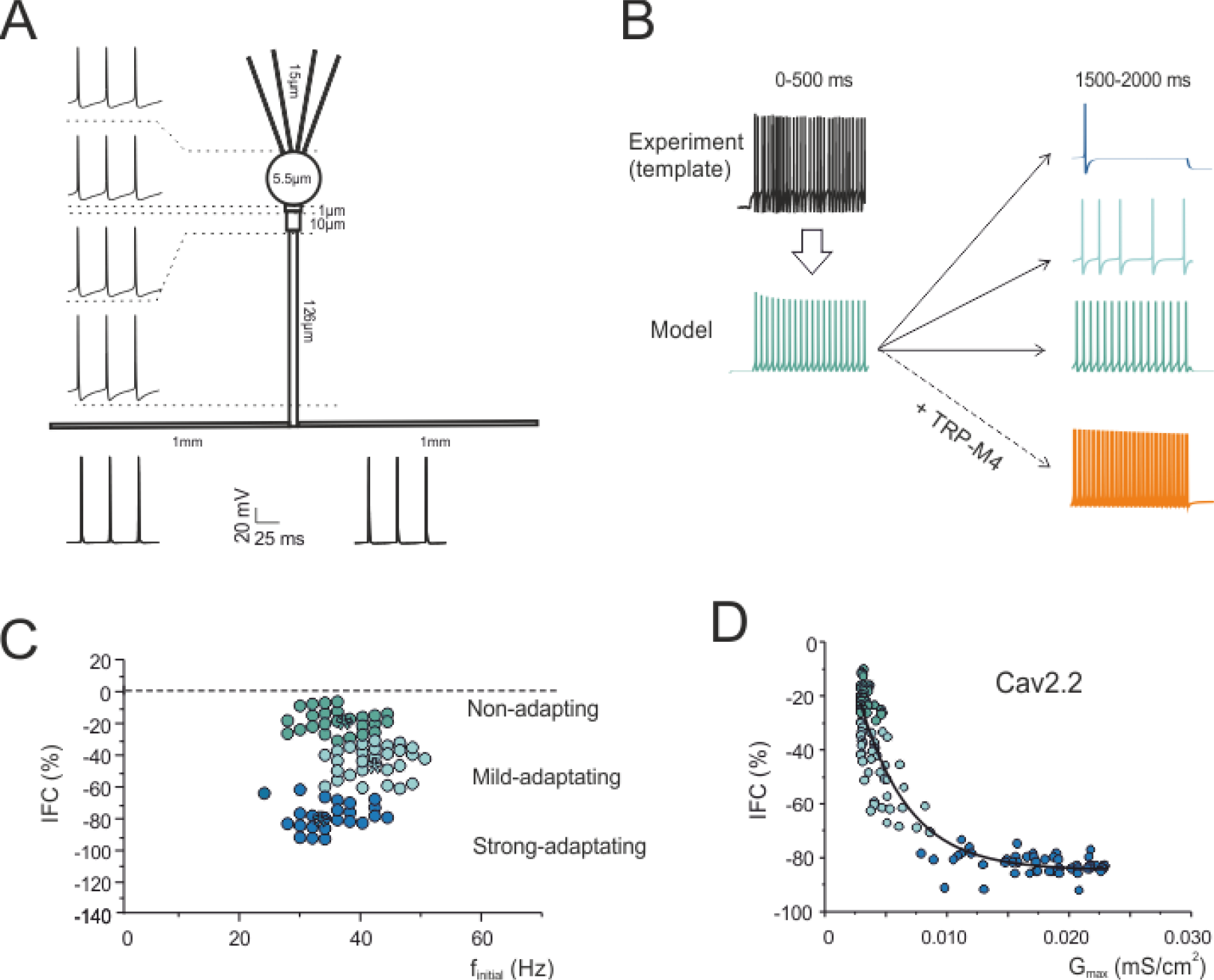
Modeling GrC and MF-GrC richness of properties. **(A)** Schematic representation of GrC morphology (not in scale; the length sections is reported) and voltage traces generated in the different model sections during 10 pA current injection. The spike shape matches that reported by (Dover et al., 2016) and its timing and amplitude remain stable across the different sections. Tables summarizing ionic channels type and distribution are reported in Supplemental Material. **(B)** An example of “template” experimental trace (obtained using a current injection of 10 pA) and of a corresponding model trace. The optimization is run over the first 500 ms of discharge (left) of a non-adapting GrC yielding a family of solutions with maximum conductances falling within the physiological range of maximum conductance values. At later times (1500-2000 ms), the discharge patterns diverges: three examples are shown (right) for a non-adapting, mild-adapting and strong-adapting solution. Addition of TRP-M4/Ca^2+^/calmoldulin subcellular mechanisms to the model yields accelerating GrCs. **(C)** The graph reports a *k*-means cluster-analysis applied to a large population of GrC models (n=150 at 10 pA) using *IFC* and *f*_*initial*_ as features. *K*-means identifies three statistically different data clusters corresponding to non-adapting, mild-adapting and strong-adapting GrCs. The centroids of the 3 subpopulations are indicated by crosses. **(D)** The graph shows the distribution of IFC in GrC models with respect to the Cav2.2 maximum conductance (mS/cm^2^), revealing a negative correlation (R^2^=0.88, n=150, p<10^−4^). At the level of GrC subpopulations, there is a clear unbalance of Cav2.2 maximum conductance (mS/cm^2^) in favor of adapting granule cells (bottom; mean values averaged over the GrC model sub-populations identified in D).

**Figure 6.**
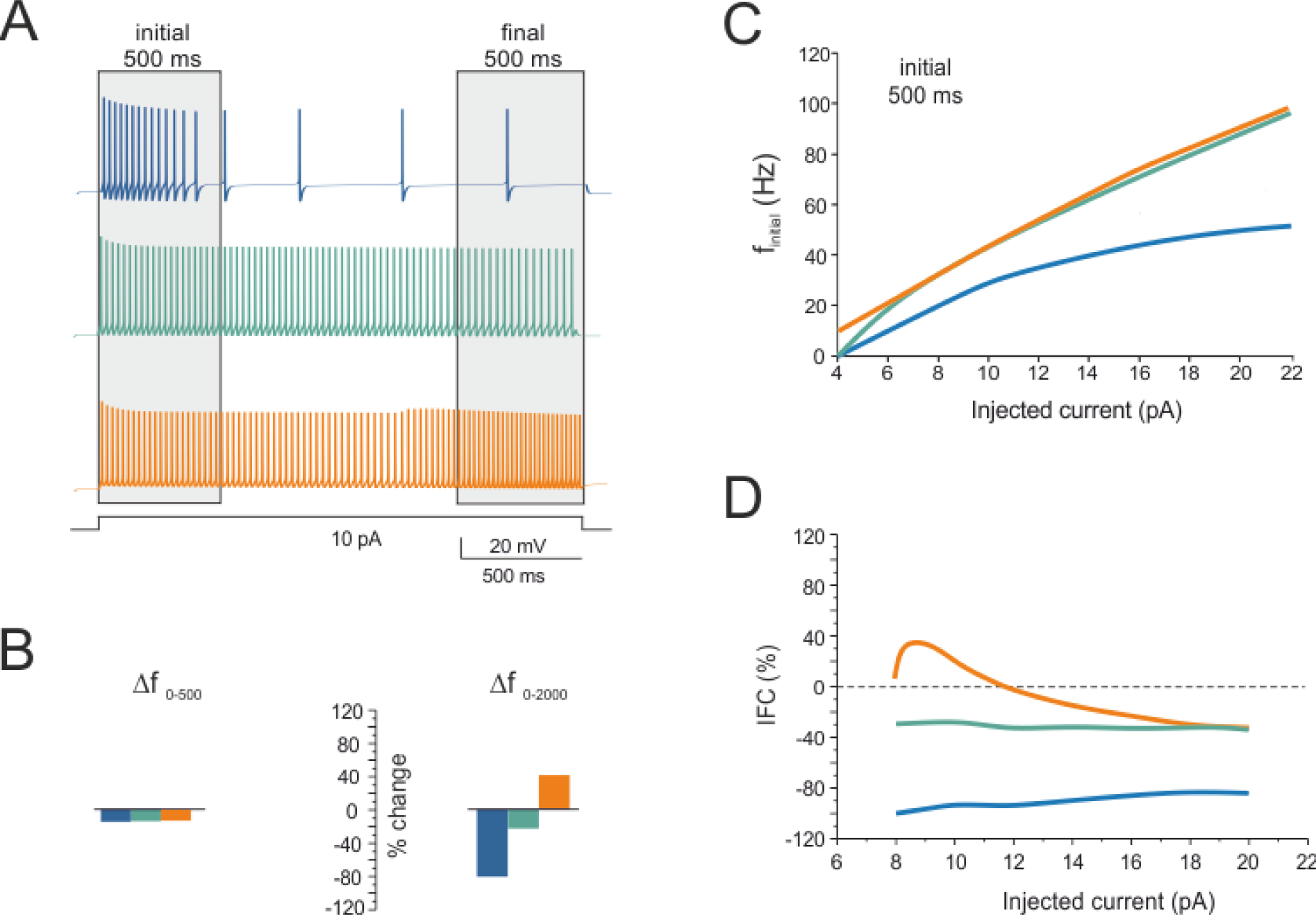
Simulation of GrC intrinsic electroresponsiveness. Three exemplar GrC model simulations are shown, one *adapting*, one *non-adapting*, one *accelerating*. The same color codes are used consistently in the figure. (A) Voltage responses to 2000 ms - 10 pA current injection from the holding potential of −65 mV. Spike frequency initially remains stable in all the three cells but it shows different trends thereafter. (B) Δf _0-500_ and Δf _0-2000_ are the spike frequency % changes after 500 ms and 2000 ms, respectively. (C) In *f*_*initial*_ /*I* plots, spike frequency increase almost linearly with the injected current intensity in both the accelerating, adapting and non-adapting granule cell models. (D) Plot of the intrinsic frequency change IFC *vs.* injected current for the three GrC models. A positive peak is apparent in the accelerating GrC at 10 pA current injection, while negative IFC values prevail in the other GrCs.

**Figure 7.**
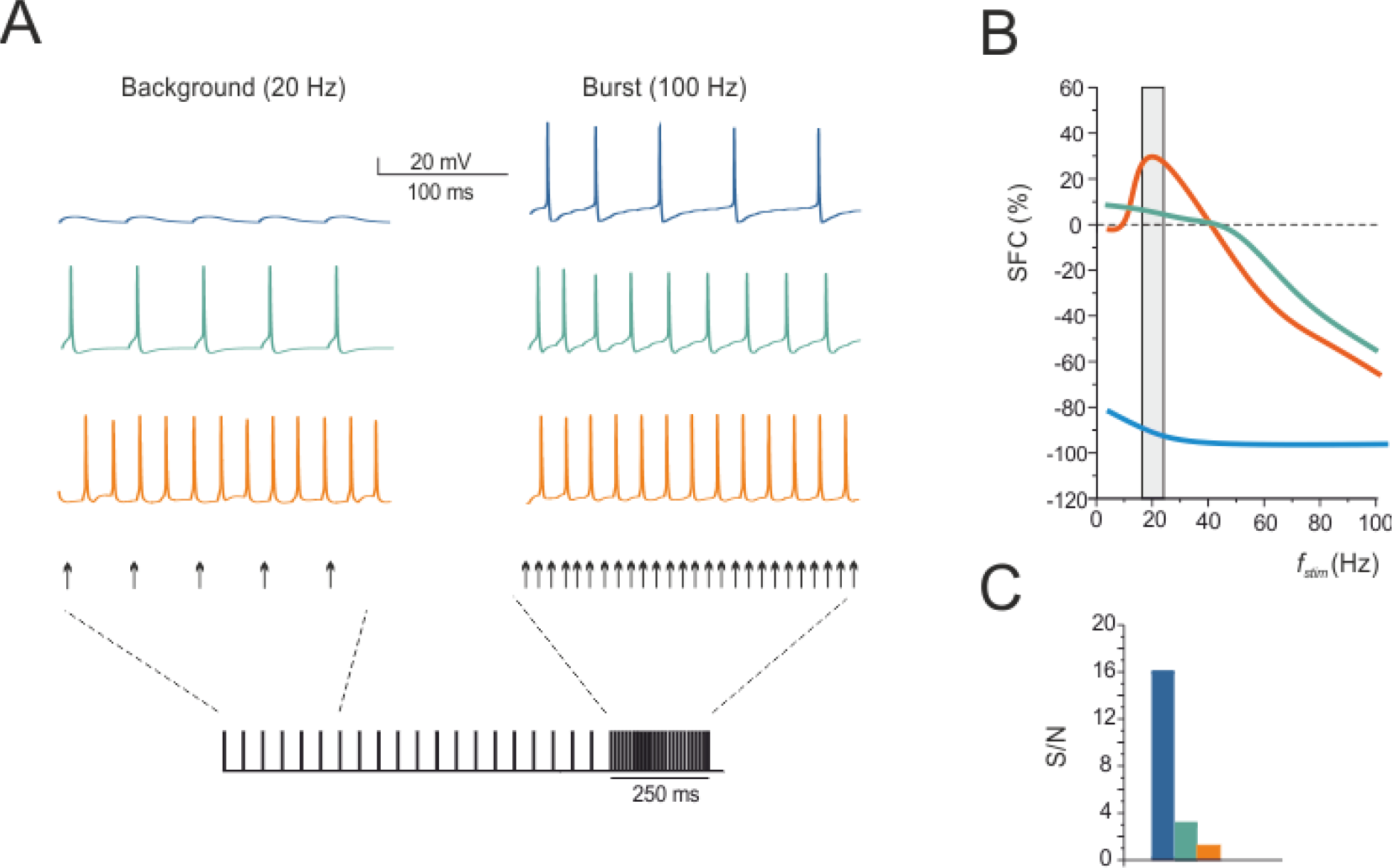
Simulation of MF-GrC synaptic transmission. Three exemplar granule cell model simulations are shown and analyzed, one *adapting*, one *non-adapting*, one *accelerating*, as defined by their intrinsic electroresponsiveness (same cells and color codes as in Fig. 6). **(A)** Voltage responses of GrC models to activation of the MF-GrC synapse (2 synapses, 1 sec continuous stimulation at 20 followed by 250 ms at 100 Hz) from the holding potential of −65 mV. The *p* values are: *adapting* GrC (*p*=0.1), *non-adapting* GrC (*p*=0.5), *accelerating* GrC (*p*=0.5) (same cells and color codes as in Fig. 6). **(B)** Plot of the synaptic frequency change SFC *vs.* stimulus frequency for the three GrC models shown in A. A positive peak was apparent in the accelerating GrC model at 20 Hz background stimulation, while negative SFC values prevailed in the other GrC models. **(C)**. Signal-to-noise ratio, (S/N = f_resp_@100 Hz/f_resp_@20 Hz) for the three GrC models in A-B. Note that S/N is much higher in adapting that in the other two GrCs.

### Mechanisms generating firing adaptation and acceleration

*Adaptation* (Fig. 8A). In the model, the larger Cav2.2 maximum conductance in strong-adapting than non-adapting GrCs (Fig. 5E) caused larger Ca^2+^ currents, which enlarged the spike upstroke by 5-10 mV, compatible with the effects of specific N-type Ca^2+^-channel blockers observed in cerebellar slices (D’Angelo et al., 1998). The larger upstroke enhanced the activation of Ca^2+^-dependent and voltage-dependent K channels, increasing K^+^ currents and spike afterhyperpolarization (AHP). The larger AHP, in turn, enhanced de-inactivation of the A-type current, which is known to protract the ISI (Connor and Stevens, 1971). Likewise, a large AHP favoured M-type current deactivation/reactivation, which can effectively slow-down (or even block) firing for hundreds of milliseconds (McCormick et al., 1992). As a whole, the model predicted that a primary increased of Ca^2+^ currents would cause a subsequent increase of K^+^ current by ~8 pA in the ISI capable of explaining firing adaptation (Fig. 8A; see also Supplemental Figs 3, 4, 5).

*Acceleration* (Fig. 8B). The TRPM4 channel, which was activated by spike trains, caused a sizeable Ca^2+^ influx through Ca^2+^ channels. Cooperative Ca^2+^ binding to Calmodulin generated a CaM2C complex that gated TRPM4 channels in a non-linear manner causing their opening once a critical concentration threshold was reached. The consequent inward current depolarized the membrane, thereby accelerating firing (see also Supplemental Fig. 3).

**Figure 8.**
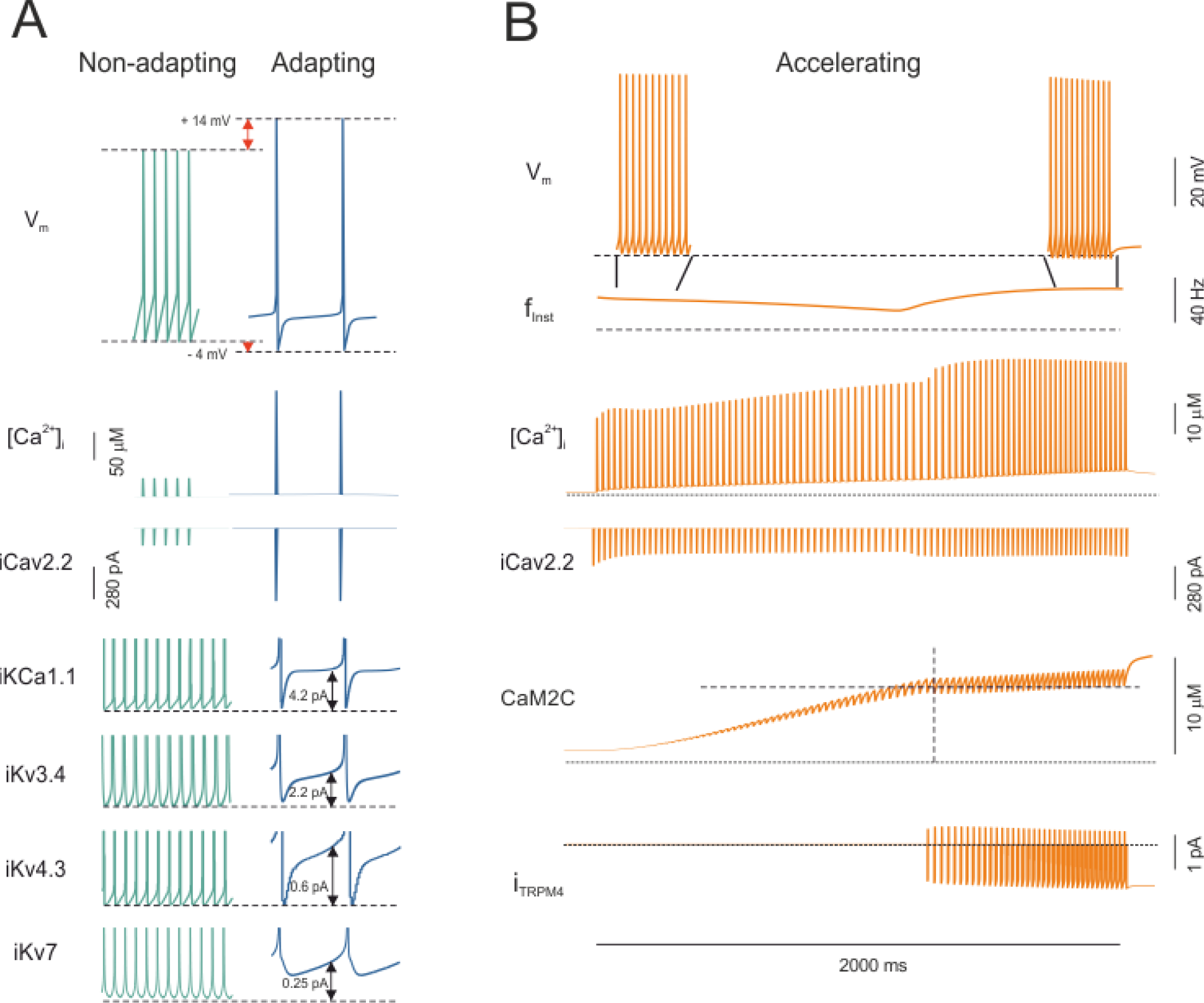
Prediction of the ionic mechanisms of firing adaptation and acceleration. **(A)** The traces show the voltage and intracellular calcium concentration along with ionic currents in non-adapting and strong-adapting GrCs during 10 pA current injection at 2000 ms. Double arrows measure the difference between non-adapting and strong-adapting GrCs. A scheme showing the hypothetical mechanisms differentiating the two cell subtypes is reported in Supplemental Material. **(B)** The traces show the voltage and intracellular calcium concentration along with ionic currents and elements of the intracellular coupling mechanism in an accelerating GrC during 10 pA current injection. A scheme showing the chemical reactions leading to TRPM4 channel opening is reported in Supplemental Material.

### Mechanisms differentiating synaptic responsiveness

EPSC trains at 100 Hz were simulated using different release probabilities (*p*=0.1, 0.5, 0.9) (Fig. 9A) and used to calculate the corresponding PPR (Sola et al., 2004). The PPR/*p* plot showed a negative slope, such that higher PPR corresponded to lower *p* values (Fig. 9B). A projection of experimental PPR values to corresponding *p* values through the PPR/*p* plot yielded *p*=0.43±0.06 for strong-adapting GrCs and to *p*=0.85±0.12 for accelerating GrCs. It should be noted that, by considering the whole PPR data distributions, *p* values in strong-adapting GrCs could range down to 0.1 and those in accelerating GrCs range up to 1.

Since *p* is a main factor regulating synaptic integration and excitation in response to input trains, the effect of different *p* values (*p*=0.1, 0.5, 0.9) was tested together with different numbers of active synapses (1-4) to systematically explore the SFC and S/N space. The mean SFC and S/N values for the four GrC groups are reported in the 3D graph of Fig. 9C and 9D. Whatever the number of active synapses, SFC at 20 Hz was larger for accelerating than strong-adapting GrCs. The low SFC and high S/N values (>10) typical of strong-adapting GrCs were found at low release probability (p=0.1), while the high SFC and low S/N values (<5) typical of all the other GrCs types were found at high release probabilities (p=0.5-0.9) in accordance to initial estimates derived from PPR analysis. Therefore, the model predicts that high S/N ratios typical of strong-adapting GrCs can be expected at low *p*, matching experimental *p* determinations.

**Figure 9.**
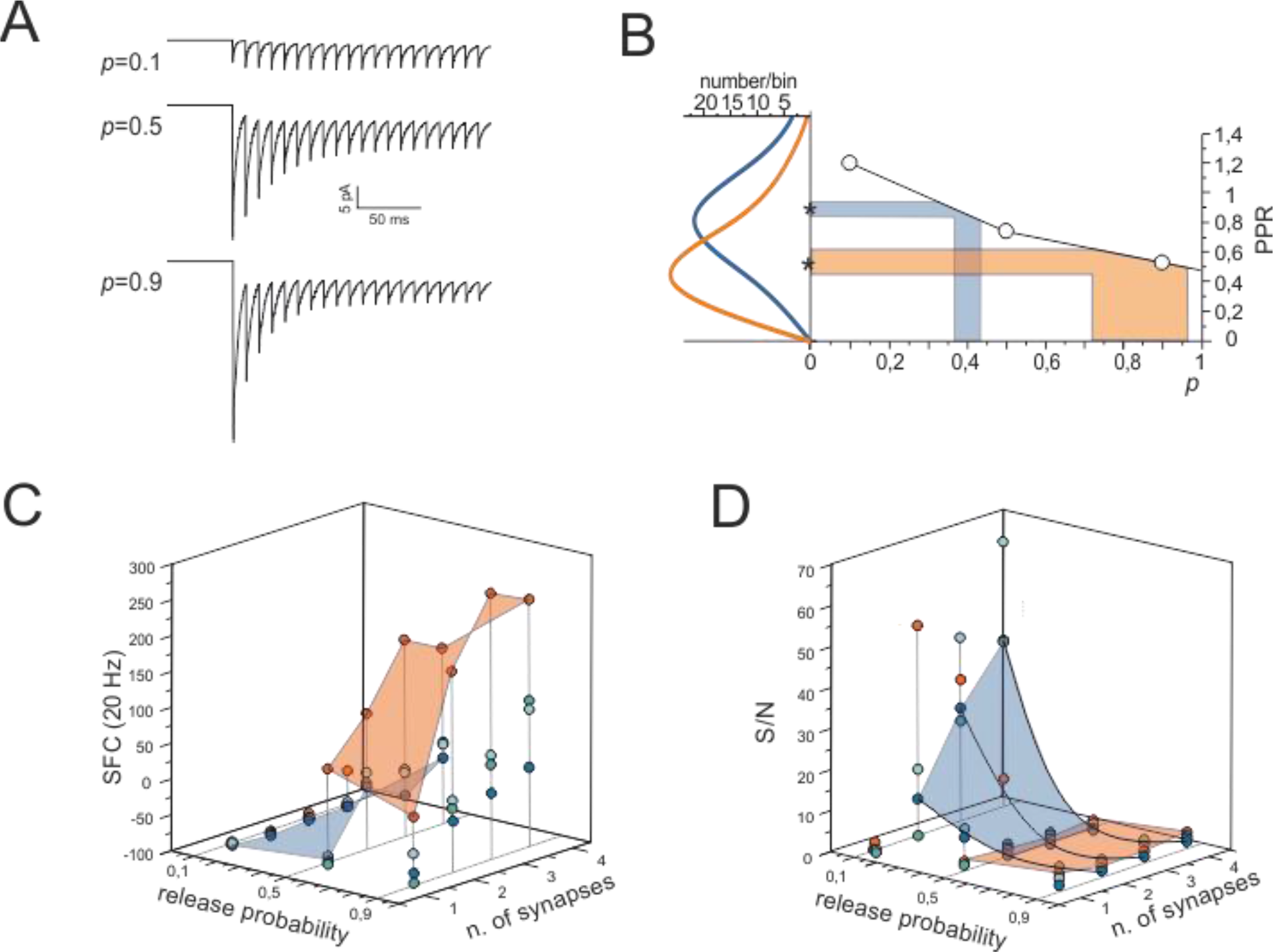
Prediction of the mechanisms of synaptic responsiveness and S/N. **(A)** The traces show voltage-clamp simulations of synaptic activity using MF-GrC synapses with a release probability *p*= 0.1, 0.5 and 0.9. Two synapses area activated in each simulations. **(B)** Prediction of *p* values from PPR data using model simulations. Stars indicate mean values of release probability in accelerating and strong-adapting granule cells and the coloured bands correspond to SEM. The probability density functions of experimental PPRs are reported on the left axis. The predicted PPR values at different *p* values are shown as open circles connected by straight lines. Through this model curve, the experimental PPRs are reflected into predicted *p* values. **(C)** Model prediction of SFC as a function of *p* and of the number of active synapses for a 20 Hz stimulation. Note that SFC is higher for accelerating than for all other GrC models at high *p*. **(D)** Model prediction of S/N (S/N = f_resp_@100 Hz/f_resp_@20 Hz) as a function of *p* and of the number of active synapses. Note that S/N is higher in strongly adapting GrC models at low *p*.

## DISCUSSION

This paper shows that cerebellar GrCs, in contrast to the canonical view describing them as a homogeneous population of neurons generating regular firing, actually show a rich repertoire of firing patterns. Over prolonged discharges (~2 sec), some GrCs remained non-adapting (20.6%) but others showed adaptation to various degree (66.7%) or, conversely, acceleration (12.7%). Adaptation and acceleration were predicted to reflect fine-tuning of membrane ionic conductances and the activation of a previously undisclosed TRPM4 channel. Specific neurotransmission properties further differentiated synaptic responsiveness.

### Are there functional granule cell subtypes?

A k-means classifier based on cerebellar GrC discharge properties uncovered four different subtypes based on intrinsic electroresposniveness: *strong-adapting*, *mild-adapting*, *non-adapting* and *accelerating* GrCs. Interestingly, these properties reverberated into response to MF stimulation (IFC/SFC plots showed a linear correlation, see Fig. 3D) suggesting that the subtypes could play a role in differentiating MF-GrC signal processing.

Since mossy fibers convey combinations of frequency modulated spike trains and bursts (Kase et al., 1980; van Kan et al., 1993; Chadderton et al., 2004; Jörntell and Ekerot, 2006; Rancz et al., 2007; Arenz et al., 2008), we evaluated how well granule cells could discriminate bursts from long-train discharges by estimating S/N. S/N was much larger in *strong-adapting* than in other GrC subtypes and this turned out to depend not only on different intrinsic electroresponsiveness but also on different release probability, *p.* According to PPR analysis (Sola et al., 2004), *p* was lower in *strong-adapting* than in the other GrC subtypes. Modeling showed indeed that, at low release probability (e.g. p=0.1-0.4), ensuing short-term facilitation can prevent low frequency background transmission, while still allowing transmission of high-frequency bursts. Conversely, at high release probability (e.g. p=0.5-0.9), ensuing short-term depression allows similar transmission of both background and bursts (cf. ((Nieus et al., 2006)). Therefore, intrinsic discharge properties and synaptic tuning concurred in differentiating the properties of synaptic excitation supporting the existence of functional MF-GrC variants specialized for differentiated signal processing.

### Parameter tuning as the basis of granule cell subtype differentiation

Parameter optimization in detailed GrC models predicted that fine tuning of membrane conductances could explain firing adaptation and acceleration.

Adaptation reflected high values of the high-threshold Ca^2+^ conductance (Rossi et al., 1994; D’Angelo et al., 1997), which increased the Ca^2+^ current and raised the spike overshoot (D’Angelo et al., 1998). This brought about a stronger activation of voltage and Ca^2+^-dependent K^+^ currents deepening the undershoot, and increasing A-type and M-type K^+^ currents. As a whole, the K^+^ current increased by ~8 pA in the ISI of strong-adapting GrCs explaining firing slowdown (GrCs have a high input resistance making the effect of a few pA current remarkable).

Acceleration was correlated with the TRPM4 current and modeling predicted that the TRPM4 channels, coupled to Calmodulin through intracellular Ca^2+^ changes, could effectively generate firing acceleration about 1.5 seconds after the beginning of discharge, i.e. when acceleration was observed experimentally (Shin et al., 2008; Shin et al., 2009; Kim et al., 2013; Menigoz et al., 2016). This delay reflected the slow cooperative gating of TRPM4 channels, that opened only after the Ca^2+^-Calmodulin complex reached a threshold.

Simulations therefore predict that fine-tuning of Ca^2+^ influx and coupling to TRPM4 channels would be critical for determining the difference between GrCs showing adaptation or acceleration.

### TRPM4 channels in granule cells

TRPM4 is a Ca^2+^-dependent non-selective cation channel, that is equally permeable to Na^+^ and K^+^, but impermeable to Ca^2+^ (Petersen, 2002). In excitable cells, it is regarded as the most suitable signaling mechanism to boost the depolarizing drive following an increase in electrical activity, which reflects into the activation of voltage-gated Ca^2+^ channels (Petersen, 2002). Emerging evidence showed that TRPM4 is actually recruited by an increase in cytosolic Ca^2+^ concentration to generate a DISC current and finely tune neuronal excitability in several brain areas (Lei et al., 2014; Menigoz et al., 2016; Riquelme et al., 2018). For instance, TRPM4 mediates the depolarizing afterpotential and phasic bursting observed in supraoptic and periventricular nuclei of the hypothalamus after a train of action potentials (Teruyama et al., 2011) and contributes to increase the firing rate in PCs (Kim et al., 2013). Here DISC TRPM4-mediated currents (blocked by the specific antagonist, 9-Phenanthrol) were first recorded in cerebellar GrCs. Interestingly, immunoistochemistry demonstrated the expression of TRPM4 in the majority of cerebellar GrCs, whereas DISC current and firing acceleration were observed in just ~12% of GrCs. It is therefore possible that the engagement of TRPM4 is finely tuned by still unknown factors. Although TRPM4 was sufficient to explain firing acceleration, we cannot exclude that other TRP channels, such as TRPM5 and TRP Canonical 5 (TRPC; e.g. see (Subramaniyam et al., 2014)) could also contribute.

### Implications for signal recoding at the cerebellum input stage

The richness of GrC intrinsic and synaptic responsiveness ends up in two main S/N patterns. In accelerating, non-adapting and mild-adapting GrCs, all frequencies are transmitted faithfully, with the accelerating GrCs being especially suitable to maintain reliable transmission at low frequencies. In strong-adapting GrCs, the low-frequencies are suppressed while the high-frequencies are transmitted, so that these neurons operate as high-pass filters. These transmission properties suggest that GrCs can process incoming MF inputs through multiple frequency-dependent filtering channels, akin with the theoretical prediction of the Adaptive Filter Model (AFM) (Dean and Porrill, 2011; Rössert et al., 2015). It is tempting to speculate that there are MF - GrC channels specialized for specific input patterns. This specialization may be the result of plasticity causing specific pre- and post-synaptic changes rewiring the system and optimizing signal transfer (e.g. LTD may characterize low-*p* MFs synapses with strong-adapting GrCs, while LTP may characterize high-*p* MFs synapses with accelerating GrCs) (Sola et al., 2004; Sgritta et al., 2017). The relationship between these putative transmission channels and zebrin stripes (Zhou et al., 2014) and GrC functional GrC clusters (Valera et al., 2016) remains to be determined.

## Conclusions

Parameter variability allowed the diversification of cerebellar granule cell firing patterns and synaptic transmission properties. Beyond that, the different balances of postsynaptic ionic conductances matched specific settings of presynaptic neurotransmitter release probability. Thus, what may simply be regarded as biological variability or noise turns out, in fact, into a richness of properties that the circuit could exploit to carry out its internal computations (Getting, 1989; Yarom and Hounsgaard, 2011; Gjorgjieva et al., 2016). Similar considerations may apply to other neurons like those of the hippocampus (Migliore et al., 2018). A question that remains to be answered is now whether differentiated neuronal and synaptic properties are determined by neuromodulatory processes or induced by plasticity. Moreover, it would be important to determine how these properties are spatially distributed inside the cerebellar circuit thereby generating specific retransmission channels shaping spatio-temporal recoding and adaptive filtering of incoming spike trains (Marr, 1969; Dean and Porrill, 2011; D’Angelo, 2016).

## Acknowledgments

We thank Giorgia Pellavio and Simona Tritto for assistance in histological procedures and Centro Grandi Strumenti. This research was supported by the European Union’s Horizon 2020 Framework Programme for Research and Innovation under Specific Grant Agreement 720270 (Human Brain Project SGA1) and 785907 (Human Brain Project SGA2). This research was supported by the HBP Neuroinformatics Platform, HBP Brain Simulation Platform, HBP HPAC Platform, funded from the European Union’s Horizon 2020 Framework Programme for Research and Innovation under the Specific Grant Agreement No. 785907 (Human Brain Project SGA2), PRACE Project 2018184373.

## REFERENCES

Arenz A, Silver RA, Schaefer AT, Margrie TW (2008) The contribution of single synapses to sensory representation in vivo. Science 321:977–980.

Arleo A, Nieus T, Bezzi M, D’Errico A, D’Angelo E, Coenen OJ (2010) How synaptic release probability shapes neuronal transmission: information-theoretic analysis in a cerebellar granule cell. Neural Comput 22:2031–2058.

Brickley SG, Cull-Candy SG, Farrant M (1996) Development of a tonic form of synaptic inhibition in rat cerebellar granule cells resulting from persistent activation of GABAA receptors. The Journal of physiology 497 (Pt 3:753–759.

Cajal SR (1911) Histologie du Système Nerveux de l’Homme et des Vertébrés, vol. II, A. Maloine Edition. Paris.

Cathala L, Brickley S, Cull-Candy S, Farrant M (2003) Maturation of EPSCs and intrinsic membrane properties enhances precision at a cerebellar synapse. The Journal of neuroscience : the official journal of the Society for Neuroscience 23:6074–6085.

Chadderton P, Margrie TW, Häusser M (2004) Integration of quanta in cerebellar granule cells during sensory processing. Nature 428:856–860.

Connor JA, Stevens CF (1971) Prediction of repetitive firing behaviour from voltage clamp data on an isolated neurone soma. J Physiol 213:31–53.

D’Angelo E, Rossi P, Taglietti V (1993) Different proportions of N-methyl-D-aspartate and non-N-methyl-D-aspartate receptor currents at the mossy fibre-granule cell synapse of developing rat cerebellum. Neuroscience 53:121–130.

D’Angelo E, De Filippi G, Rossi P, Taglietti V (1995) Synaptic excitation of individual rat cerebellar granule cells in situ: evidence for the role of NMDA receptors. The Journal of physiology 484 (Pt 2:397–413.

D’Angelo E, De Filippi G, Rossi P, Taglietti V (1997) Synaptic activation of Ca2+ action potentials in immature rat cerebellar granule cells in situ. J Neurophysiol 78:1631–1642.

D’Angelo E, De Filippi G, Rossi P, Taglietti V (1998) Ionic mechanism of electroresponsiveness in cerebellar granule cells implicates the action of a persistent sodium current. J Neurophysiol 80:493–503.

D’Angelo E, Rossi P, Armano S, Taglietti V (1999) Evidence for NMDA and mGlu receptor-dependent long-term potentiation of mossy fiber-granule cell transmission in rat cerebellum. Journal of neurophysiology 81:277–287.

D’Angelo E, Nieus T, Maffei a, Armano S, Rossi P, Taglietti V, Fontana a, Naldi G (2001) Theta-frequency bursting and resonance in cerebellar granule cells: experimental evidence and modeling of a slow k+-dependent mechanism. The Journal of neuroscience : the official journal of the Society for Neuroscience 21:759–770.

D’Errico A, Prestori F, D’Angelo E (2009) Differential induction of bidirectional long-term changes in neurotransmitter release by frequency-coded patterns at the cerebellar input. J Physiol 587:5843–5857.

Dean P, Porrill J (2011) Evaluating the adaptive-filter model of the cerebellum. J Physiol 589:3459–3470.

Dean P, Porrill J (2014) Decorrelation learning in the cerebellum: computational analysis and experimental questions. Prog Brain Res 210:157–192.

Deb K, Pratap A, Agarwal S, Meyarivan T (2002) A fast and elitist multiobjective genetic algorithm: NSGA-II. IEEE Transactions on Evolutionary Computation 6:182–197.

Diwakar S, Magistretti J, Goldfarb M, Naldi G, D’Angelo E (2009) Axonal Na+ channels ensure fast spike activation and back-propagation in cerebellar granule cells. Journal of neurophysiology 101:519–532.

Dover K, Marra C, Solinas S, Popovic M, Subramaniyam S, Zecevic D, D’Angelo E, Goldfarb M (2016) FHF-independent conduction of action potentials along the leak-resistant cerebellar granule cell axon. Nature communications 7:12895–12895.

Druckmann S, Banitt Y, Gidon A, Schürmann F, Markram H, Segev I (2007) A novel multiple objective optimization framework for constraining conductance-based neuron models by experimental data. Frontiers in neuroscience 1:7–18.

Druckmann S, Berger TK, Schürmann F, Hill S, Markram H, Segev I (2011) Effective stimuli for constructing reliable neuron models. PLoS computational biology 7:e1002133–e1002133.

D’Angelo E (2016) Challenging Marr’s theory of the cerebellum. In, pp 62–78: Oxford University Press.

Fuhrmann G, Segev I, Markram H, Tsodyks M (2002) Coding of temporal information by activity-dependent synapses. J Neurophysiol 87:140–148.

Gall D, Roussel C, Susa I, D’Angelo E, Rossi P, Bearzatto B, Galas MC, Blum D, Schurmans S, Schiffmann SN (2003) Altered neuronal excitability in cerebellar granule cells of mice lacking calretinin. The Journal of neuroscience : the official journal of the Society for Neuroscience 23:9320–9327.

Getting PA (1989) Emerging principles governing the operation of neural networks. Annu Rev Neurosci 12:185–204.

Gjorgjieva J, Drion G, Marder E (2016) Computational implications of biophysical diversity and multiple timescales in neurons and synapses for circuit performance. Curr Opin Neurobiol 37:44–52.

Goldfarb M, Schoorlemmer J, Williams A, Diwakar S, Wang Q, Huang X, Giza J, Tchetchik D, Kelley K, Vega A, Matthews G, Rossi P, Ornitz DM, D’Angelo E (2007) Fibroblast Growth Factor Homologous Factors Control Neuronal Excitability through Modulation of Voltage-Gated Sodium Channels. Neuron 55:449–463.

Golgi C (1906) The neuron doctrine-theory and facts. Nobel Lectures: Physiology or Medicine:29–29.

Heath NC, Rizwan AP, Engbers JDT, Anderson D, Zamponi GW, Turner RW (2014) The Expression Pattern of a Cav3-Kv4 Complex Differentially Regulates Spike Output in Cerebellar Granule Cells. Journal of Neuroscience 34:8800–8812.

Herculano-Houzel S (2010) Coordinated scaling of cortical and cerebellar numbers of neurons. Frontiers in neuroanatomy 4:12.

Hines ML, Carnevale NT (2001) Neuron: A Tool for Neuroscientists. The Neuroscientist 7:123–135.

Hines ML, Carnevale NT (2008) Translating network models to parallel hardware in NEURON. Journal of neuroscience methods 169:425–455.

Hines ML, Davison AP, Muller E (2009) NEURON and Python. Frontiers in neuroinformatics 3:1–1.

Houston CM, Diamanti E, Diamantaki M, Kutsarova E, Cook A, Sultan F, Brickley SG (2017) Exploring the significance of morphological diversity for cerebellar granule cell excitability. Scientific reports 7:46147–46147.

Jörntell H, Ekerot CF (2006) Properties of somatosensory synaptic integration in cerebellar granule cells in vivo. J Neurosci 26:11786–11797.

Kase M, Miller DC, Noda H (1980) Discharges of Purkinje cells and mossy fibres in the cerebellar vermis of the monkey during saccadic eye movements and fixation. J Physiol 300:539–555.

Kim YS, Kang E, Makino Y, Park S, Shin JH, Song H, Launay P, Linden DJ (2013) Characterizing the conductance underlying depolarization-induced slow current in cerebellar Purkinje cells. Journal of Neurophysiology.

Laforenza U, Scaffino MF, Gastaldi G (2013) Aquaporin-10 represents an alternative pathway for glycerol efflux from human adipocytes. PLoS One 8:e54474.

Lei YT, Thuault SJ, Launay P, Margolskee RF, Kandel ER, Siegelbaum SA (2014) Differential contribution of TRPM4 and TRPM5 nonselective cation channels to the slow afterdepolarization in mouse prefrontal cortex neurons. Front Cell Neurosci 8:267.

Mapelli L, Gagliano G, Soda T, Laforenza U, Moccia F, D’Angelo EU (2017) Granular Layer Neurons Control Cerebellar Neurovascular Coupling Through an NMDA Receptor/NO-Dependent System. J Neurosci 37:1340–1351.

Marr BYD (1969) A theory of cerebellar cortex. J Physiol 202:437–470.

Masoli S, Rizza MF, Sgritta M, Van Geit W, Schürmann F, D’Angelo E (2017) Single Neuron Optimization as a Basis for Accurate Biophysical Modeling: The Case of Cerebellar Granule Cells. Frontiers in Cellular Neuroscience 11:1–14.

McCormick DA, Strowbridge BW, Huguenard J (1992) Determination of State-Dependent Processing in Thalamus by Single Neuron Properties and Neuromodulators. In, pp 259–290: Elsevier.

Menigoz A, Ahmed T, Sabanov V, Philippaert K, Pinto S, Kerselaers S, Segal A, Freichel M, Voets T, Nilius B, Vennekens R, Balschun D (2016) TRPM4-dependent post-synaptic depolarization is essential for the induction of NMDA receptor-dependent LTP in CA1 hippocampal neurons. Pflugers Arch 468:593–607.

Migliore R et al. (2018) The physiological variability of channel density in hippocampal CA1 pyramidal cells and interneurons explored using a unified data-driven modeling workflow. PLoS Comput Biol 14:e1006423.

Mitchell SJ, Silver RA (2000a) Glutamate spillover suppresses inhibition by activating presynaptic mGluRs. Nature 404:498–502.

Mitchell SJ, Silver RA (2000b) GABA spillover from single inhibitory axons suppresses low-frequency excitatory transmission at the cerebellar glomerulus. J Neurosci 20:8651–8658.

Mrejeru A, Wei A, Ramirez JM (2011) Calcium-activated non-selective cation currents are involved in generation of tonic and bursting activity in dopamine neurons of the substantia nigra pars compacta. J Physiol 589:2497–2514.

Nieus T, Sola E, Mapelli J, Saftenku E, Rossi P, D’Angelo E (2006) LTP regulates burst initiation and frequency at mossy fiber-granule cell synapses of rat cerebellum: experimental observations and theoretical predictions. Journal of neurophysiology 95:686–699.

Nieus TR, Mapelli L, D’Angelo E (2016) Corrigendum: Regulation of output spike patterns by phasic inhibition in cerebellar granule cells. Front Cell Neurosci 10:30.

Nilius B, Prenen J, Tang J, Wang C, Owsianik G, Janssens A, Voets T, Zhu MX (2005) Regulation of the Ca2+ sensitivity of the nonselective cation channel TRPM4. J Biol Chem 280:6423–6433.

Petersen OH (2002) Cation channels: homing in on the elusive CAN channels. Curr Biol 12:R520–522.

Rancz Ea, Ishikawa T, Duguid I, Chadderton P, Mahon S, Häusser M (2007) High-fidelity transmission of sensory information by single cerebellar mossy fibre boutons. Nature 450:1245–1248.

Riquelme D, Silva I, Philp AM, Huidobro-Toro JP, Cerda O, Trimmer JS, Leiva-Salcedo E (2018) Subcellular Localization and Activity of TRPM4 in Medial Prefrontal Cortex Layer 2/3. Front Cell Neurosci 12:12.

Rossi P, D’Angelo E, Magistretti J, Toselli M, Taglietti V (1994) Age-dependent expression of high-voltage activated calcium currents during cerebellar granule cell development in situ. Pflugers Arch 429:107–116.

Rossi P, De Filippi G, Armano S, Taglietti V, D’Angelo E (1998) The weaver mutation causes a loss of inward rectifier current regulation in premigratory granule cells of the mouse cerebellum. J Neurosci 18:3537–3547.

Rössert C, Dean P, Porrill J (2015) At the Edge of Chaos: How Cerebellar Granular Layer Network Dynamics Can Provide the Basis for Temporal Filters. PLoS Comput Biol 11:e1004515.

Saviane C, Silver RA (2006) Fast vesicle reloading and a large pool sustain high bandwidth transmission at a central synapse. Nature 439:983–987.

Sgritta M, Locatelli F, Soda T, Prestori F, D’Angelo EU (2017) Hebbian Spike-Timing Dependent Plasticity at the Cerebellar Input Stage. J Neurosci 37:2809–2823.

Shin JH, Kim YS, Linden DJ (2008) Dendritic glutamate release produces autocrine activation of mGluR1 in cerebellar Purkinje cells. Proc Natl Acad Sci U S A 105:746–750.

Shin JH, Kim YS, Worley PF, Linden DJ (2009) Depolarization-induced slow current in cerebellar Purkinje cells does not require metabotropic glutamate receptor 1. Neuroscience 162:688–693.

Silver RA, Traynelis SF, Cull-Candy SG (1992) Rapid-time-course miniature and evoked excitatory currents at cerebellar synapses in situ. Nature 355:163–166.

Silver RA, Colquhoun D, Cull-Candy SG, Edmonds B (1996) Deactivation and desensitization of non-NMDA receptors in patches and the time course of EPSCs in rat cerebellar granule cells. J Physiol 493 (Pt 1):167–173.

Sola E, Prestori F, Rossi P, Taglietti V, D’Angelo E (2004) Increased neurotransmitter release during long-term potentiation at mossy fibre-granule cell synapses in rat cerebellum. The Journal of Physiology 557:843–861.

Solinas S, Nieus T, D’Angelo E (2010) A realistic large-scale model of the cerebellum granular layer predicts circuit spatio-temporal filtering properties. Frontiers in cellular neuroscience 4:12–12.

Stokum JA, Kwon MS, Woo SK, Tsymbalyuk O, Vennekens R, Gerzanich V, Simard JM (2018) SUR1-TRPM4 and AQP4 form a heteromultimeric complex that amplifies ion/water osmotic coupling and drives astrocyte swelling. Glia 66:108–125.

Subramaniyam S, Solinas S, Perin P, Locatelli F, Masetto S, D’Angelo E (2014) Computational modeling predicts the ionic mechanism of late-onset responses in unipolar brush cells. Front Cell Neurosci 8:237.

Teruyama R, Sakuraba M, Kurotaki H, Armstrong WE (2011) Transient receptor potential channel m4 and m5 in magnocellular cells in rat supraoptic and paraventricular nuclei. J Neuroendocrinol 23:1204–1213.

Tsodyks M, Pawelzik K, Markram H (1998) Neural networks with dynamic synapses. Neural Comput 10:821–835.

Valera AM, Binda F, Pawlowski SA, Dupont JL, Casella JF, Rothstein JD, Poulain B, Isope P (2016) Stereotyped spatial patterns of functional synaptic connectivity in the cerebellar cortex. Elife 5.

Van Geit W (2015) Blue Brain Project (2015). eFEL.Available online at: https://github.com/BlueBrain/eFEL (Accessed February 16, 2016). In.

Van Geit W, Gevaert M, Chindemi G, Rössert C, Courcol J-D, Muller EB, Schürmann F, Segev I, Markram H (2016) BluePyOpt: Leveraging Open Source Software and Cloud Infrastructure to Optimise Model Parameters in Neuroscience. Frontiers in Neuroinformatics 10:1–30.

van Kan PL, Gibson AR, Houk JC (1993) Movement-related inputs to intermediate cerebellum of the monkey. J Neurophysiol 69:74–94.

Yarom Y, Hounsgaard J (2011) Voltage fluctuations in neurons: signal or noise? Physiol Rev 91:917–929.

Zhou H, Lin Z, Voges K, Ju C, Gao Z, Bosman LW, Ruigrok TJ, Hoebeek FE, De Zeeuw CI, Schonewille M (2014) Cerebellar modules operate at different frequencies. Elife 3:e02536.

Zitzler E, Künzli S (2004) Indicator-Based Selection in Multiobjective Search. In, pp 832–842.

