## Supplemental materials for "Parameter tuning differentiates granule cell subtypes enriching the repertoire of retransmission properties at the cerebellum input stage"

### SUPPLEMENTAL MATERIAL

#### *Ionic channels*

*Nav1.6* – The spike generation and propagation mechanisms exploited different sodium channels in the axonal hillock and initial segment (Nav1.6-FHF) with respect to AA and PF (Nav1.6) (Khaliq et al., 2003; Khaliq and Raman, 2006; Magistretti et al., 2006; Goldfarb et al., 2007; Dover et al., 2010; Dover et al., 2016). Therefore we developed two versions of the Nav1.6 sodium channel: 1) without FHF in AA and PF, and 2) with FHF in Hillock and AIS (Goldfarb et al., 2007). The channel without FHF was taken from (Diwakar et al., 2009a). The channel with FHF was based (Dover et al., 2010) and modified with a faster kinetic for the Loff transition (from 0.15/ms to 0.5/ms). The FHF channel in the AIS had a higher conductance compared to channels in the other section, in order to compensate for the absence of somatic sodium channels.

*Kv1.1, Kv1.5, Kv2, Kv3.4, Kv4.3, Kv7/Km* – To counterbalance the resurgent current of both types of sodium currents, the axonal sections were endowed with the Kv3.4 potassium channel (Khaliq et al., 2003; Chang et al., 2007; Masoli et al., 2015). The A-type potassium current (Kv4.3) was taken from the canonical mono and multi compartmental models (D'Angelo et al., 2001; Diwakar et al., 2009a) and placed on the soma. To reduce the calcium excitability (Khavandgar et al., 2005), the potassium channel Kv1.1 was placed, alone, on the dendrites and mixed with Kv1.5 on the soma (Chung et al., 2001). The potassium current Kv2, taken from Channelpedia (http://channelpedia.epfl.ch/ionchannels/193), was placed on the soma to act as slow delayed rectifying channel (Debanne et al., 2011; Ranjan et al., 2011). The GrC M-current (Kv7), taken from the mono compartmental model (D'Angelo et al., 2001) was placed on the AIS, instead of the soma, in accordance with its *anchoring system which linked it to* Ankyrin-G enriched zones, usually found in the AIS (Cooper, 2011).

*Cav2.2, KCa1.1* – The GrC N-type high-voltage activated calcium channel, Cav2.2, was taken from the previous mono compartmental model (D'Angelo et al., 2001) and distributed along the entire morphology. The main calcium dependent potassium channels, Kca1.1 was placed only on the dendrites.

*Kir2.x* – The GrC K inward rectifier channel (Rossi et al., 2006) was taken from (D'Angelo et al., 2001; Diwakar et al., 2009a) and placed in the soma.

TRPM - A TRPM - like current was taken from the UBC multicompartmental model (Subramaniyam et al., 2014), modified to interact with Calmodulin (Pepke et al., 2010) in the 2C conformation (Cam2C) (Nilius et al., 2005) when reached a certain threshold. This channel was placed only on the dendrites.

*Calcium ($Ca^{2+}$) buffer* - GrC are known to buffer intracellular calcium through Calretinin (Gall et al., 2003; Schwaller, 2014). To account for this, the $Ca^{2+}$ buffer previously used by (Masoli et al., 2015; Masoli et al., 2017) by changing Calbindin and Parvalbumin with the 12 states kinetic Calretinin kinetic model (Faas et al., 2007). The basal concentration of the protein was set accordingly to experimental data (Saftenku, 2012).

***Supplemental Table 1. Electrotonic compartments in the GrC model.***

| *Section name* | *Diameter (μm)* | *Length (μm)* | *N° of sections* | *Cm* (μF/cm$^2$) | *Leak Rp* | *Leak Gmax* (S/cm$^2$) |
|---|---|---|---|---|---|---|
| **Dendrites** | 0.75 | 15 | 4 | 2.5 | -60 | 0.0002 - 0.0004 |
| **Soma** | 5.8 | 5.62 | 1 | 2 | -60 | 0.0002 - 0.0004 |
| **Hillock** | 1.5 | 1 | 1 | 2 | -60 | 0.0002 - 0.0004 |
| **AIS** | 0.7 | 10 | 1 | 1 | -60 | 0.0002 - 0.0004 |
| **AA** | 0.3 | 126 | 18 | 1 | -60 | 0.0002 - 0.0004 |
| **PF** | 0.15 | 980 | 140 | 1 | -60 | 0. 0000001 - 0.0000008 |

The table shows the sections of the GrC model along with their number, diameter and length. GrC model dendrites, soma and hillock were taken unmodified from (Masoli et al., 2017), whereas the morphology was extended to accommodate the Axon Initial Segment (AIS) along with a specific mechanism for spike generation (Bender and Trussell, 2012). The AIS was built as a single section 10μm long and 0.7μm wide (Debanne et al., 2011; Dover et al., 2016). The ascending axon (AA) and parallel fibers (PF) were built connecting 7μm modules until reaching the desired axonal length (Wilms and Häusser, 2015). The present model consisted of an AA composed by 18 sections (total length of 126μm) and two PF branches composed of 140 sections (1mm each). The diameters were set to 0.3 μm for AA and 0.15 μm for PF (Dover et al., 2016). Membrane capacitance ($C_m$) was set at 1 μF/cm$^2$ in the AIS, AA and PFs, 2.5 μF/cm$^2$ in the dendrites and 2 μF/cm$^2$ in the soma, hillock (these higher $C_m$ values were used to normalize spike amplitude). The axial resistance ($R_a$) was set at 100 **Ω***cm in the entire model. The input resistance ($R_{in}$), calculated from a 10mV current transient (from -70 to -80 mV), in voltage-clamp mode, was in the range of 1.2-1.8 GΩ (e.g. (D'Angelo et al., 1995)).

***Supplemental Table 2. Ionic mechanisms in the GrC model.***

| Conductance/ Location | | Range $G_{i\text{-}max}$ ($S/cm^2$) | $E_{rev}$ (mV) | Description of channel (H.H or Markovian) | Reference |
|---|---|---|---|---|---|
| **Na channels** | | | | | |
| Nav1.6 FHF | Hillock | 0.008 - 0.05 | 87.39 | Markovian | (Magistretti et al., 2006; Diwakar et al., 2009b) |
| | AIS | 1 – 1.65 | | | |
| Nav1.6 No FHF | aa | 0.02 - 0.05 | | | |
| | pf | 0.01 - 0.04 | | | |
| **K channels** | | | | | |
| Kv1.1 | Dendrites | 0.0001 - 0.01 | -88 | HH | (Akemann and Knopfel, 2006) |
| | Soma | 0.003 – 0.01 | | | |
| Kv1.5 | Soma | 0.13e-4 - 0.13e-2 | -88 | HH | (Courtemanche et al., 1998) |
| Kv2 | Soma | 0.000001 - 0.0001 | -88 | HH | (Ranjan et al., 2011) |
| Kv3.4 | Soma | 0.0005 - 0.005 | -88 | HH | (Raman and Bean, 2001; Khaliq et al., 2003) |
| | Hillock | 0.02 - 0.08 | | | |
| | AIS | 0.005 - 0.04 | | | |
| | aa | 0.002 - 0.005 | | | |
| | pf | 0.004 - 0.01 | | | |
| Kv4.3 | Soma | 0.002 - 0.004 | -88 | HH | (D'Angelo et al., 2001; Diwakar et al., 2009b) |
| Km | AIS | 0.0003 - 0.0008 | -88 | HH | (D'Angelo et al., 2001; Diwakar et al., 2009b) |
| Kir2.x | Soma | 0.0005 - 0.001 | -88 | HH | (D'Angelo et al., 2001; Diwakar et al., 2009b) |
| **Ca dependent K channels** | | | | | |
| Kca1.1 | Dendrites | 0.010 - 0.03 | -88 | Markovian | (Anwar et al., 2010) |
| **Ca channels** | | | | | |
| Cav2.2 | Dendrites | 0.005 - 0.025 | 137.5 | HH | (D'Angelo et al., 2001; Diwakar et al., 2009b) |
| | Soma | 0.0001 - 0.0007 | | | |
| | Hillock | 0.0001 - 0.0007 | | | |
| | AIS | 0.0001 - 0.0007 | | | |
| | aa | 0.0001 - 0.0007 | | | |
| | pf | | | | |
| **TRPM4 like channel** | | | | | |
| TRPM4 | Dendrites | $5*10^{-4}$ | 0 | HH | (Subramaniyam et al., 2014) |
| **Calcium buffer - Pumps density** | | | | | |
| Ca Buffer | Dendrites | $1*10^{-9}$ | | Markovian | Based on (Anwar et al., 2010) Modified with data from (Faas et al., 2007; Saftenku, 2012) |
| | Soma | $1*10^{-9}$ | | | |
| | Hillock | $1*10^{-9}$ | | | |
| | AIS | $1*10^{-9}$ | | | |
| | aa | $1*10^{-9}$ | | | |
| | pf | | | | |

The table reports the ionic channels, their location, the range of the maximum conductances and the ionic reversal potential. The corresponding gating equations were written either in Hodgkin-Huxley (HH) style or in Markovian style according to the indicated references.

***Supplemental Table 3. Spike features.***

| | ***10 pA*** | ***16 pA*** | ***22 pA*** |
|---|---|---|---|
| | ***Exp*** | ***Exp*** | ***Exp*** |
| Spike height (mV) | 20.93 | 19.25 | 18.88 |
| Spike width (mV) | 0.70 | 1.0 | 1.06 |
| AHP depth (mV) | -71.24 | -61.85 | -57.25 |
| AHP depth Slow (mV) | -57.93 | -52.90 | -51.52 |
| Time to first spike (ms) | 121.29 | 107.42 | 107.10 |
| Mean spike frequency (Hz) | 63.54 | 97.28 | 98.90 |
| ISI CV | 0.18 | 0.0983 | 0.088 |
| Spike Count | 141 | 202 | 207 |

The table shows exemplar values of features, obtained from experimental traces in a granule cell using eFEL.

### SUPPLEMENTAL FIGURES

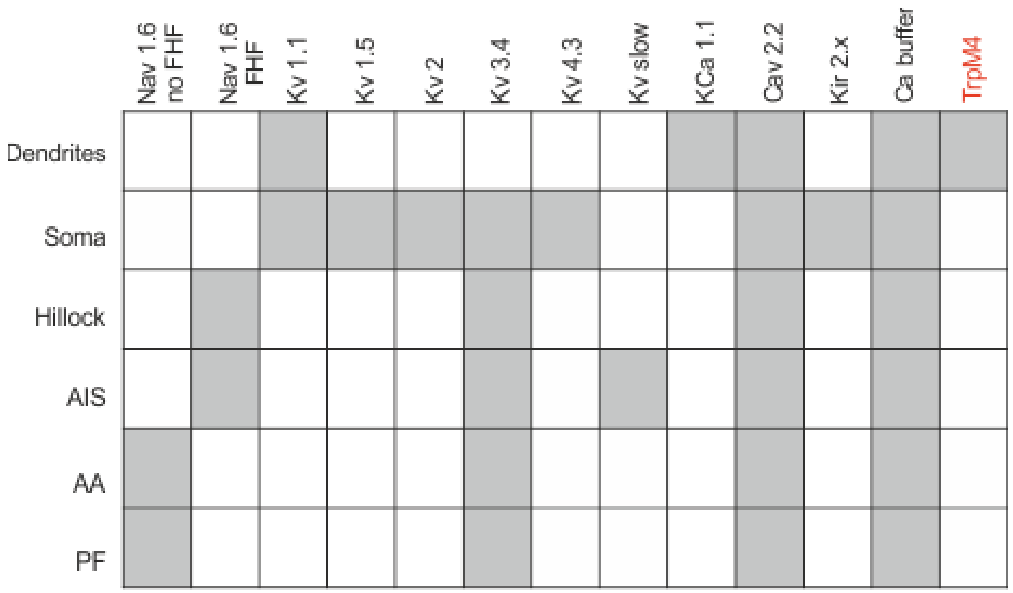


**Figure S1.** ***Distribution of ionic channels in the granule cell model.*** The table summarizes the ionic channel type and distribution in the different GrC model sections according to literature. The description, for each ionic channel, can be found in the previous section of the supplemental material.

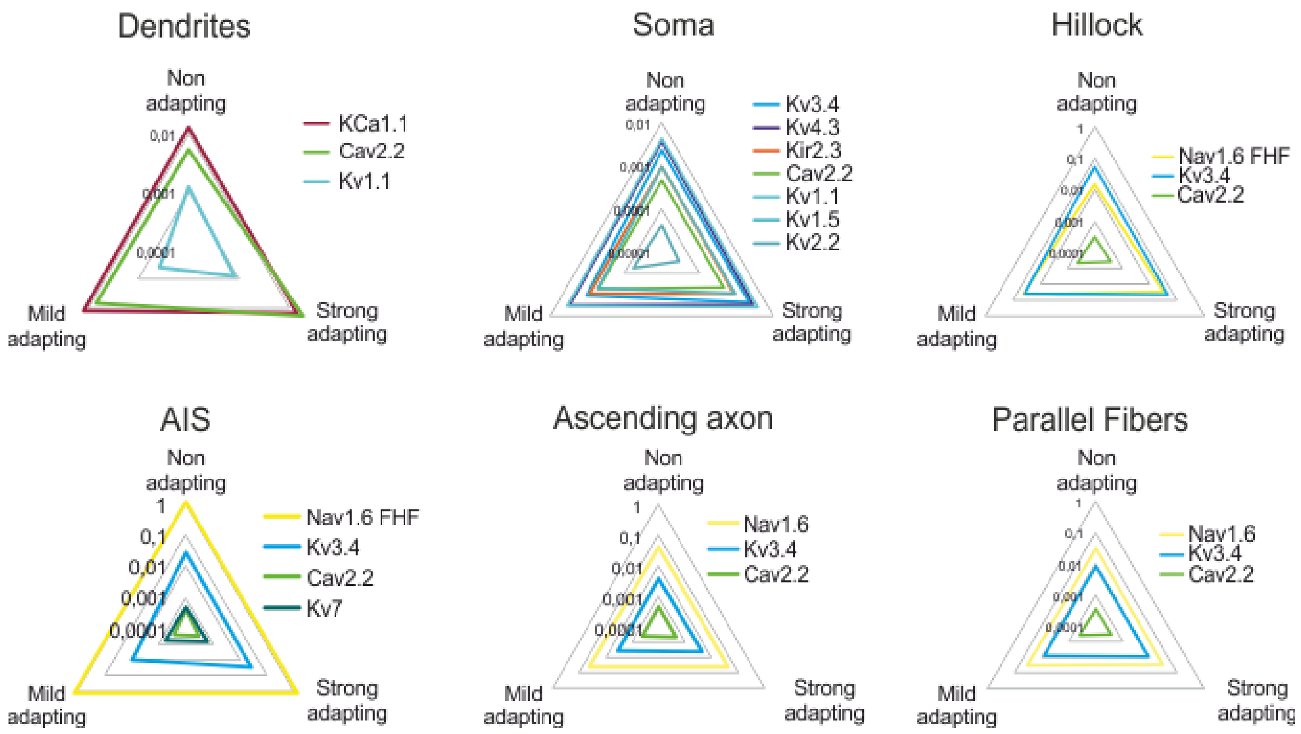


**Figure S2.** ***Computational modelling of granule cell electroresponsiveness***. (A) Comparison of average conductance values in the three GrC subtypes. The conductances were obtained from the optimization and describe the differences in each section along the morphology. Notably, strong-adapting granule cells had a markedly higher Cav2.2, slightly higher Nav1.6-FHF and Kv3.4 compared to the other granule cells.

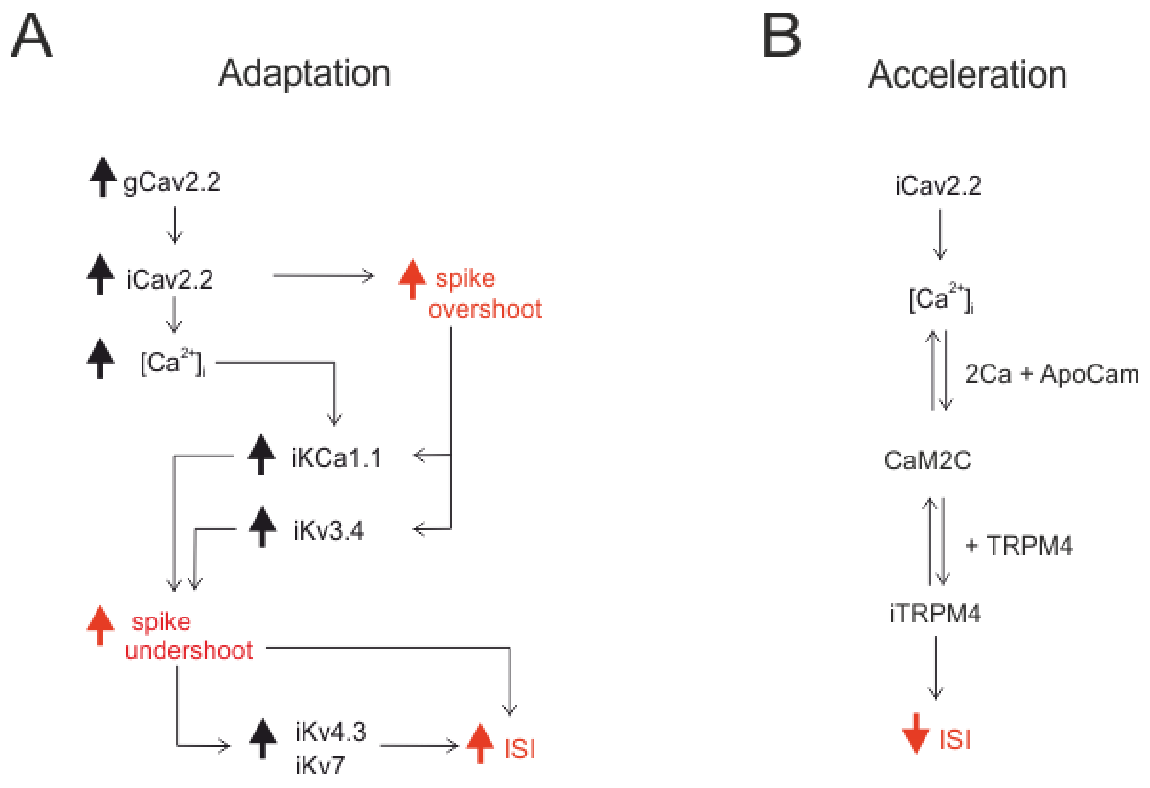


***Figure S3. Hypothesis on the mechanisms of adaptation and acceleration.*** (A) *Adaptation.* Schematics of the hypothetic chain of events leading to adaptation. The chain is started by increased $Ca^{2+}$ conductance. (B) *Acceleration.* Schematics of the hypothetic steps required for firing acceleration. The critical step is coupling of $Ca^{2+}$ influx to Calmodulin leading to TRPM4 channel opening.

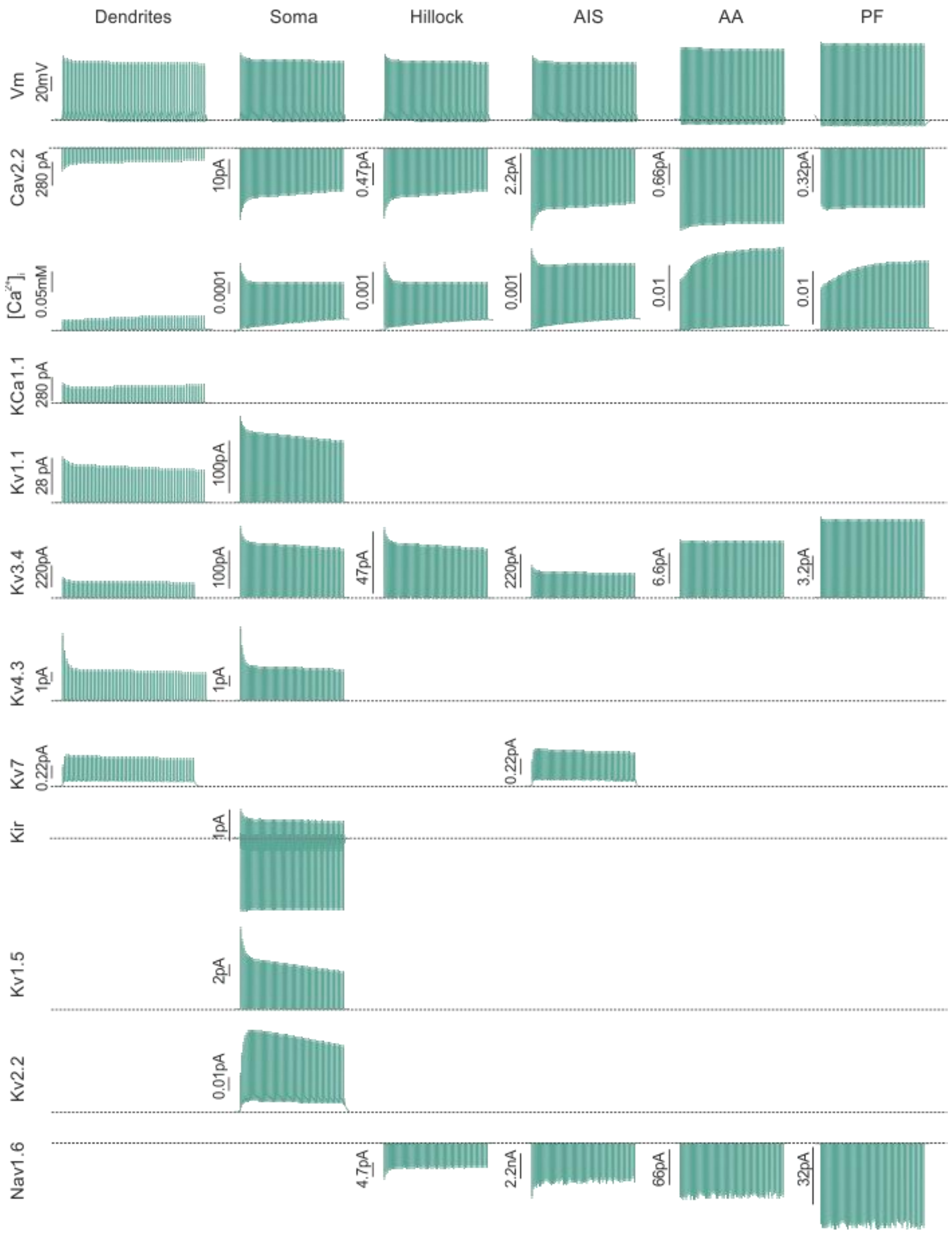


***Figure S4. Ionic currents in non-adapting granule cells during firing.*** The currents, for each ionic channel, were recorded from a dendrite, the soma, hillock, Axon Initial Segment, a distal section of AA and a distal section of PF.

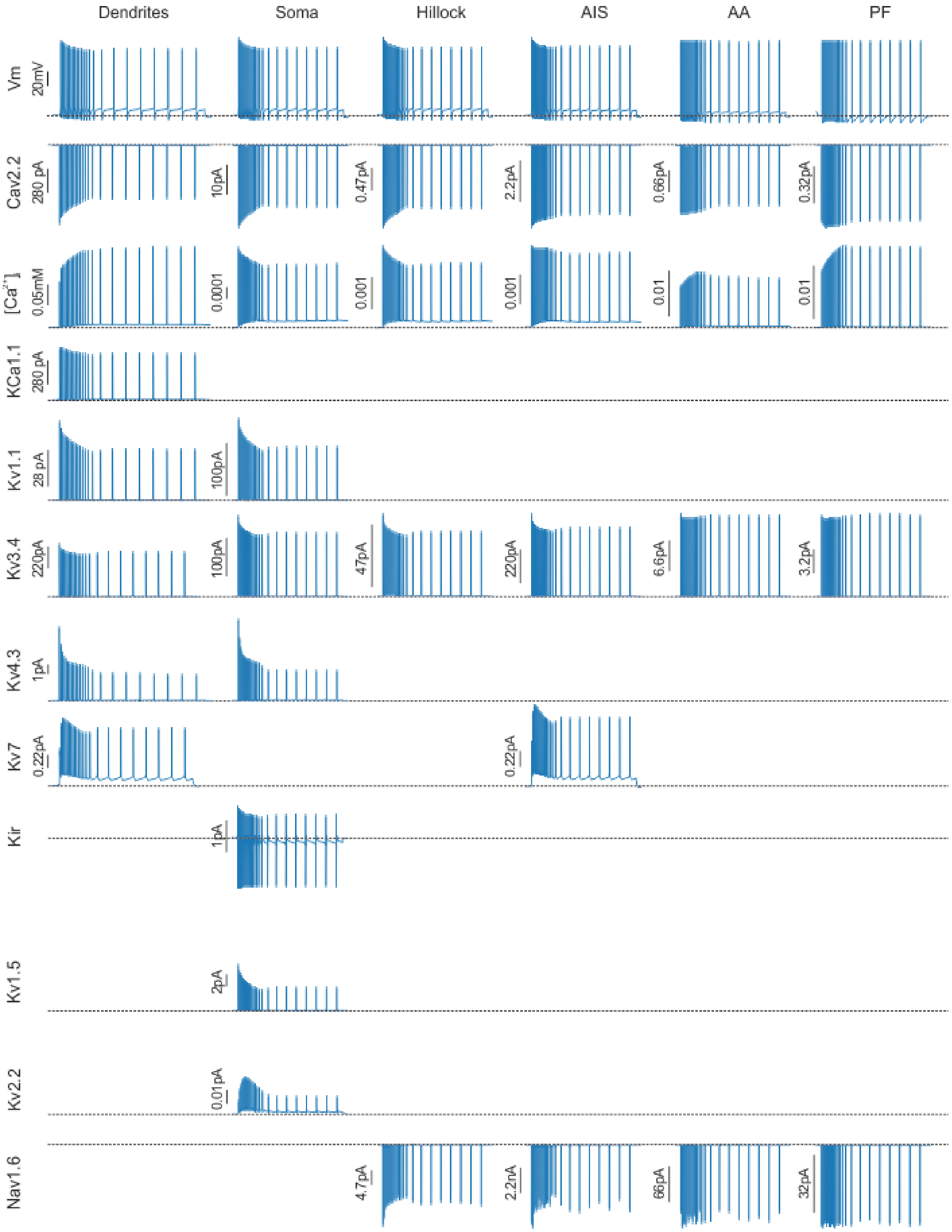


***Figure S5. Ionic currents in strongly adapting granule cells during firing.*** The currents, for each ionic channel, were recorded from a dendrite, the soma, hillock, Axon Initial Segment, a distal section of AA and a distal section of PF.

### REFERENCES


Akemann W, Knopfel T (2006) Interaction of Kv3 Potassium Channels and Resurgent Sodium Current Influences the Rate of Spontaneous Firing of Purkinje Neurons. Channels 26:4602-4612.

Anwar H, Hong S, De Schutter E (2010) Controlling Ca2+-Activated K+ Channels with Models of Ca2+ Buffering in Purkinje Cells. The Cerebellum:1-13.

Bender KJ, Trussell LO (2012) The Physiology of the Axon Initial Segment. Annual Review of Neuroscience 35:249-265.

Chang SY, Zagha E, Kwon ES (2007) Distribution of Kv3.3 potassium channel subunits in distinct neuronal populations of mouse brain. Journal of comparative neurology 972:953-972.

Chung YH, Shin C, Kim MJ, Lee BK, Cha CI (2001) Immunohistochemical study on the distribution of six members of the Kv1 channel subunits in the rat cerebellum. Brain Res 895:173-177.

Cooper EC (2011) Made for "anchorin": Kv7.2/7.3 (KCNQ2/KCNQ3) channels and the modulation of neuronal excitability in vertebrate axons. Seminars in Cell & Developmental Biology 22:185-192.

Courtemanche M, Ramirez RJ, Nattel S (1998) Ionic mechanisms underlying human atrial action potential properties: insights from a mathematical model. The American journal of physiology 275:H301-321.

D'Angelo E, De Filippi G, Rossi P, Taglietti V (1995) Synaptic excitation of individual rat cerebellar granule cells in situ: evidence for the role of NMDA receptors. The Journal of physiology 484 ( Pt 2:397-413.

D'Angelo E, Nieus T, Maffei a, Armano S, Rossi P, Taglietti V, Fontana a, Naldi G (2001) Theta-frequency bursting and resonance in cerebellar granule cells: experimental evidence and modeling of a slow k+-dependent mechanism. The Journal of neuroscience : the official journal of the Society for Neuroscience 21:759-770.

Debanne D, Campanac E, Bialowas A, Carlier E, Alcaraz G (2011) Axon physiology. Physiol Rev 91:555-602.

Diwakar S, Magistretti J, Goldfarb M, Naldi G, D'Angelo E (2009a) Axonal Na+ channels ensure fast spike activation and back-propagation in cerebellar granule cells. Journal of neurophysiology 101:519-532.

Diwakar S, Magistretti J, Goldfarb M, Naldi G, D'Angelo E (2009b) Axonal Na+ channels ensure fast spike activation and back-propagation in cerebellar granule cells. Journal of neurophysiology 101:519-532.

Dover K, Solinas S, D'Angelo E, Goldfarb M (2010) Long-term inactivation particle for voltage-gated sodium channels. The Journal of physiology 588:3695-3711.

Dover K, Marra C, Solinas S, Popovic M, Subramaniyam S, Zecevic D, D'Angelo E, Goldfarb M (2016) FHF-independent conduction of action potentials along the leak-resistant cerebellar granule cell axon. Nature communications 7:12895-12895.

Faas GC, Schwaller B, Vergara JL, Mody I (2007) Resolving the fast kinetics of cooperative binding: Ca2+ buffering by calretinin. PLoS Biology 5:2646-2660.

Gall D, Roussel C, Susa I, D'Angelo E, Rossi P, Bearzatto B, Galas MC, Blum D, Schurmans S, Schiffmann SN (2003) Altered neuronal excitability in cerebellar granule cells of mice lacking calretinin. The Journal of neuroscience : the official journal of the Society for Neuroscience 23:9320-9327.

Goldfarb M, Schoorlemmer J, Williams A, Diwakar S, Wang Q, Huang X, Giza J, Tchetchik D, Kelley K, Vega A, Matthews G, Rossi P, Ornitz DM, D'Angelo E (2007) Fibroblast Growth Factor Homologous Factors Control Neuronal Excitability through Modulation of Voltage-Gated Sodium Channels. Neuron 55:449-463.

Khaliq ZM, Raman IM (2006) Relative contributions of axonal and somatic Na channels to action

potential initiation in cerebellar Purkinje neurons. Journal of Neuroscience 26:1935-1944.
Khaliq ZM, Gouwens NW, Raman IM (2003) The contribution of resurgent sodium current to high-frequency firing in Purkinje neurons: an experimental and modeling study. Journal of Neuroscience 23:4899-4912.
Khavandgar S, Walter JT, Sageser K, Khodakhah K (2005) Kv1 channels selectively prevent dendritic hyperexcitability in rat Purkinje cells. The Journal of physiology 569:545-557.
Magistretti J, Castelli L, Forti L, D'Angelo E (2006) Kinetic and functional analysis of transient, persistent and resurgent sodium currents in rat cerebellar granule cells in situ: an electrophysiological and modelling study. The Journal of physiology 573:83-106.
Masoli S, Sergio S, Egidio DA (2015) Action potential processing in a detailed Purkinje cell model reveals a critical role for axonal compartmentalization. Frontiers in Cellular Neuroscience 9:1--22.
Masoli S, Rizza MF, Sgritta M, Van Geit W, Schürmann F, D'Angelo E (2017) Single Neuron Optimization as a Basis for Accurate Biophysical Modeling: The Case of Cerebellar Granule Cells. Frontiers in Cellular Neuroscience 11:1-14.
Nilius B, Prenen J, Tang J, Wang C, Owsianik G, Janssens A, Voets T, Zhu MX (2005) Regulation of the Ca2+ sensitivity of the nonselective cation channel TRPM4. J Biol Chem 280:6423-6433.
Pepke S, Kinzer-Ursem T, Mihalas S, Kennedy MB (2010) A dynamic model of interactions of Ca2+, calmodulin, and catalytic subunits of Ca2+/calmodulin-dependent protein kinase II. PLoS Comput Biol 6:e1000675.
Raman IM, Bean BP (2001) Inactivation and recovery of sodium currents in cerebellar Purkinje neurons: evidence for two mechanisms. Biophysical Journal 80:729-737.
Ranjan R, Khazen G, Gambazzi L, Ramaswamy S, Hill SL, Schürmann F, Markram H (2011) Channelpedia: An Integrative and Interactive Database for Ion Channels. Frontiers in Neuroinformatics 5:1-8.
Rossi P, Mapelli L, Roggeri L, Gall D, De Kerchove D'Exaerde A, Schiffmann SN, Taglietti V, D'Angelo E (2006) Inhibition of constitutive inward rectifier currents in cerebellar granule cells by pharmacological and synaptic activation of GABAB receptors. European Journal of Neuroscience 24:419-432.
Saftenku EÈ (2012) Effects of Calretinin on Ca2+ Signals in Cerebellar Granule Cells: Implications of Cooperative Ca2+ Binding. The Cerebellum 11:102-120.
Schwaller B (2014) Calretinin: from a "simple" Ca(2+) buffer to a multifunctional protein implicated in many biological processes. Frontiers in neuroanatomy 8:3-3.
Subramaniyam S, Solinas S, Perin P, Locatelli F, Masetto S, D'Angelo E (2014) Computational modeling predicts the ionic mechanism of late-onset responses in unipolar brush cells. Front Cell Neurosci 8:237.
Wilms CD, Häusser M (2015) Reading out a spatiotemporal population code by imaging neighbouring parallel fibre axons in vivo. Nature Communications 6:6464-6464.
